# Regulation of PL fimbriae (*plf*) and role of PlfB as a negative regulator during avian pathogenic *Escherichia coli* infection in turkeys

**DOI:** 10.64898/2026.10.07.757495

**Authors:** Fariba Akrami, Tylor Pereira, Sébastien Houle, Hossein Jamali, Evelyne Ng Kwan Lim, Eric Massé, France Daigle, Charles M. Dozois

## Abstract

Little is currently known about the regulation of PL fimbriae produced by some pathogenic *E. coli* strains. We identified PlfB, a transcriptional regulator of the PapB family, as a negative regulator of PL fimbriae (Plf). Deletion of *plfB* significantly increased *plfA* transcription and Plf production. PlfB bound specifically to the *plfA* promoter region. Using a P*plfA*–*lux* reporter system, the deletion of *plfB* caused sustained and increased expression across serial passages. Global regulatory genes such as *lrp*, *argR*, and *hns* were also implicated in regulation of *plf* expression. Loss of *lrp* or *argR* decreased *plf* expression, suggesting these regulators positively regulate PL fimbriae. By contrast, loss of *hns* increased levels of *plf* expression, supporting its role as a negative regulator. Interestingly, loss of *plfB* significantly increased bacterial colonization in the lung of 6-day-old turkey poults. Further, complementation of *plfB* restored *plf* fimbrial expression and colonization of turkey lungs to wild-type levels. This further supports an important role for PL fimbriae for colonization of turkey lungs, since loss of *plfB* which resulted in increased *plf* gene expression in turkey lungs also correlated with higher bacterial numbers in the lungs. Taken together, these results indicate that PlfB acts as a negative regulator to reduce levels of PL fimbrial expression, under some conditions, but that it is not required for *plf* expression. This contrasts with the role the PapB regulator, which has been shown to play an essential role in expression of P fimbriae in uropathogenic *E. coli* strains.

**IMPORTANCE:** Regulation of *E. coli* PL fimbriae, encoded by *plf*, remains poorly understood. We investigated the role of a novel PapB-family transcriptional regulator, PlfB. In a *plfB* mutant *plf* transcription and fimbrial production were increased. In gel shift assays PlfB bound to the *plfA* promoter region. A P*plfA*–*lux* reporter further demonstrated increased and sustained expression in the *plfB* mutant. Deletion of genes encoding global regulators supported positive roles for Lrp and ArgR and a negative role for H-NS in *plf* regulation. Importantly, deletion of *plfB* significantly increased colonization of turkey lungs. These findings identify PlfB as a direct negative regulator of PL fimbriae and demonstrate that modulation of *plf* expression influences bacterial colonization during avian infection. Importantly, in contrast to most other PapB-family regulators, PlfB was shown to play a negative role in regulation of its cognate fimbrial system and was not required for optimal expression of fimbriae.

## Introduction

Most *Escherichia coli* strains are commensal bacteria. However, some strains are pathogenic and can cause either intestinal or extraintestinal infections in humans and other animals, such as avian pathogenic *E. coli* (APEC) (1). The pathogenic potential of *E*. *coli* strains is influenced by the presence of different virulence genes, which are located on the bacterial chromosome or on some types of plasmids, such as IncFIB/IncFIC virulence plasmids. One group of these plasmids comprise the ColV-like plasmids that are present in most APEC strains (2–5). Recently, a new type of fimbria (pilus) called P-like (PL) fimbria encoded by the *plf* genes was identified on a ColV-type plasmid (pEC598) from APEC strain QT598, and these *plf* genes were also found to be present in a variety of *E*. *coli* strains isolated from poultry, humans, and other sources (6).

Fimbrial adhesins, as a key class of virulence factors, are proteinaceous structures assembled on the surface of bacteria to contribute to biofilm formation, bacterial attachment and colonization and can contribute to evasion of the host immune response and adaptation to environmental conditions (1, 3, 7–9). Pathogenic *E. coli* can express multiple types of fimbrial adhesins providing an adaptive advantage, by enabling bacterial cells to recognize distinct host cell receptors and increase host-range and tissue tropisms (6). The type 1 and P (also called Pap) fimbriae have been studied extensively and are commonly produced by extraintestinal pathogenic *E. coli,* such as APEC as well as human extra-intestinal *E*. *coli* (ExPEC) (3, 7, 10–12). P fimbriae and other related systems belong to the π fimbrial family, and also include PL, Pix, Sfp, and Pef fimbriae (6, 13).

Transcriptional regulation, phase variation, DNA methylation, and small RNAs are types of regulatory mechanisms allowing bacterial populations to switch from expression one type of fimbriae to another in response to environmental changes or stresses, including temperature, osmolarity, nutrient status, and host signals (14–16). Fimbrial systems are tightly regulated at the transcriptional and post-transcriptional levels by global regulators (for example H-NS, Lrp, and CRP) and specific dedicated regulators (such as FimB and Fim E recombinases for regulation of type 1 fimbriae (17) and PapB and related regulators that mediate regulation of P fimbriae and a variety of other fimbrial systems (18). These regulatory mechanisms allow bacterial population adaptability through expression of multiple types of fimbriae in response to environmental cues and signals within different host niches (1, 3, 7–9).

Among the global regulators involved in fimbrial control, H-NS generally acts as a transcriptional silencer and contributes to environmentally responsive regulation of both type 1 and P fimbriae, whereas Lrp participates directly in the phase-variable regulation of these fimbrial systems (11, 17). CRP-cAMP further links fimbrial expression to the metabolic state of the cell and can influence both type 1 fimbrial phase switching and transcriptional control of the *pap* operon (11, 16). In the *pap* system, Lrp and the local regulator PapI cooperate to establish alternative methylation dependent ON and OFF states, with PapI promoting Lrp binding to sites associated with the transcriptionally active state. PapB provides an additional level of control by regulating *papI* and *papBA* transcription and can also modulate expression of other fimbrial systems, highlighting the broader role of PapB-family regulators in coordinating fimbrial expression (3, 16).

PL fimbriae mediate biofilm production, hemagglutination, and bacterial attachment to human bladder and kidney epithelial cells, contribute to kidney colonization in a murine UTI model (6) and to extraintestinal infection of turkey poults in a respiratory infection model (18) The *plf* gene cluster comprises at least 10 predicted genes, and includes a predicted member of the PapB family, PlfB, which shares 45% identity/ 67% similarity to the PapB regulator (6). However, unlike the P fimbrial system, the Plf system lacks any equivalent of the PapI regulator (6). Since there is little information concerning mechanisms of regulation of PL fimbriae, the aim of this work was to investigate mechanisms of regulation of PL fimbriae including the potential role of the PlfB gene and to identify the environmental and genetic factors controlling PL fimbrial expression. Herein, we demonstrate that PL fimbriae are also affected by global regulators such as Lrp and H-NS. Further, we demonstrate that PlfB seems to act as a negative regulator of PL fimbrial expression, as loss of this gene results in increased expression of PL fimbriae and increased colonization of turkey lungs following infection.

## Results

### Evaluation of *plf* expression *in vitro* using a *plf* promoter *lux* fusion

The PL fimbrial system comprises 9 genes corresponding to predicted structural proteins, and a predicted regulatory protein that shares some identity to the PapB regulatory protein, which we have called PlfB. However, regulation of expression of these fimbriae have not been investigated. The PL fimbriae revealed a similarity to the Pap family with a notable divergence in the regulatory region and a distinct type of G adhesin (Fig. 1A) (6). We predicted putative promoter regions for investigating the transcriptional regulation by the upstream intergenic region (http://www.softberry.com/cgi-bin/programs/gfindb/bprom.pl) (Fig. 1B). The 6 predicted promoters were suggested based on predicted −35/−10 motifs and presence of 2 open reading frames of putative genes that could encode regulators, which we have named *plfX* and *plfB,* and a predicted promoter adjacent to the *plfA* gene which encodes the major fimbrial subunit.

**FIG. 1.**
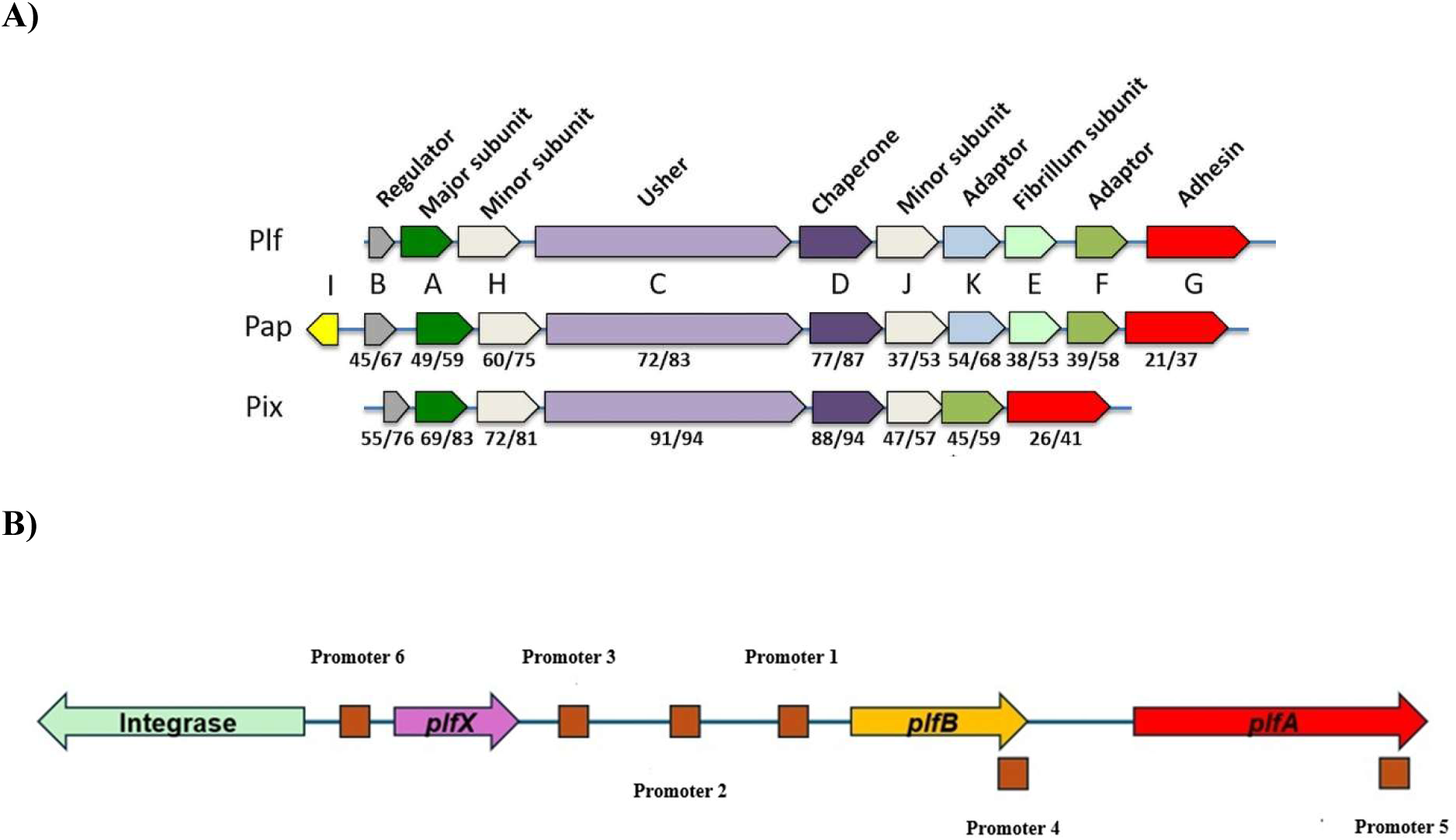
PL fimbrial (*plf*) gene cluster comparison and predicted promoters. **A)** Comparative genetic organization of the *plf, pap*, and *pix* fimbrial gene clusters. Gene arrangements are shown with conserved functional classes annotated (regulator, major subunit, minor subunits, chaperone, usher, adaptor, fibrillum subunit, and adhesin) (6) Numbers below genes for the *pap* and *pix* gene clusters indicate the % identity/% similarity of the proteins to the corresponding genes of the Plf system. (B) Six putative promoter regions (P1–P6) were identified within the regulatory region based on predicted −35 and −10 motifs. These included predicted promoters upstream of *plfB* and *plfX* and a promoter adjacent to *plfA*. These initial designations were used until promoter activity and transcriptional organization were experimentally characterized.

While *plfB* is predicted to encode a PapB-like regulatory protein, *plfX* would encode a small protein of unknown function. Given its location adjacent to the fimbrial gene cluster, *plfX* was considered as a potential regulator of *plf* transcription and was therefore also included for further analyses. These included a primary promoter immediately adjacent to *plfB* and several additional upstream regions associated with neighbouring ORFs and an integrase gene. The six putative promoter regions were initially designated P1-P6 (Fig. 1B) based on their relative positions within the regulatory region, prior to experimental validation of promoter activity. The promoter region, from promoter 6 (P6) to the end of promoter 4 (P4), was cloned into the digested Tn7-cm-Lux vector to construct pIJ610 and then transferred into MGN-617 (QT5814). Finally, *Pplf-lux* was conjugated at the Tn7 site of the QT598 (QT5828) and QT598Δ*plf* (QT5995) to investigate the promoter expression in various conditions. Analyzing the expression of the promoter region in various nutritional and environmental conditions revealed that *plf* expression was significantly upregulated after 3 passages on LBA and 0.3M NaCl at 37°C among all of those mentioned conditions in the methods (Fig. 2A). This salt concentration was selected after testing both lower and higher NaCl levels, which indicated that 0.3 M NaCl produced the most consistent increase in promoter activity while still supporting normal bacterial growth. Standard LB agar contains approximately 0.17 M NaCl, and supplementation to 0.3 M therefore represents an ∼1.8-fold increase in salt concentration. The activity of the promoter region tested following different conditions is shown in Fig. S1.

**FIG. 2.**
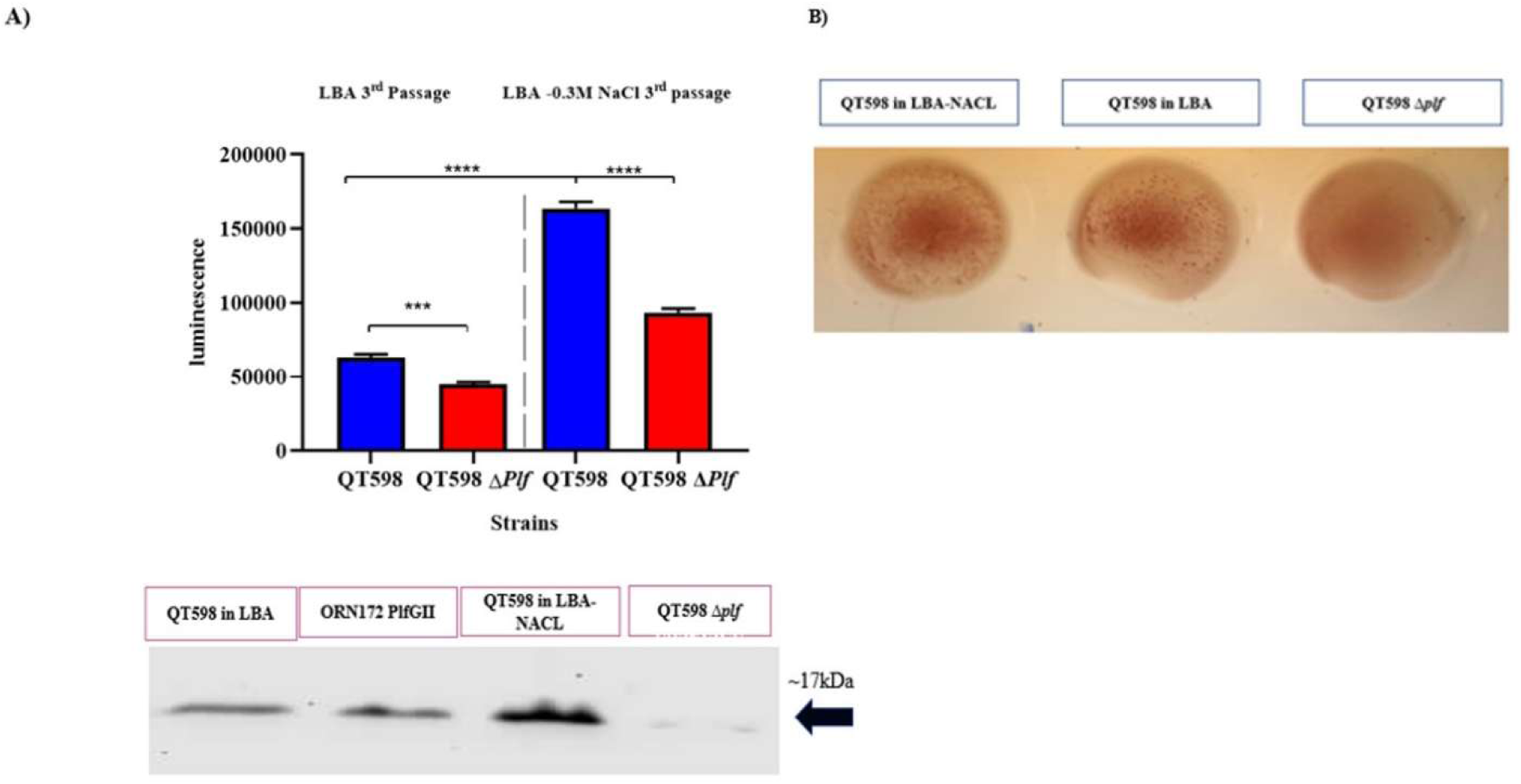
Comparative expression of *plf* after growth on LBA or LBA-0.3 M NaCl determined by *lux* reporter, western blot and hemagglutination assay. **A)** The *plf* promoter region was cloned into a mini-Tn7 suicide vector, a promoterless lux reporter, located in a mini-Tn7 suicide vector, and subsequently transferred into QT598 and QT598Δ*plf* through conjugation from *E. coli* MGN-617, which carried the delivery plasmid. Transposition was directed by the Tn7 machinery in the recipient, resulting in precise insertion of the construct at the *att*Tn7 site located immediately downstream of glmS. B) Luminescence from QT598 P*plf-lux* and QT598Δ*plf* P*plf-lux* was measured after three consecutive passages of growth on LBA and LBA-0.3M NaCl. The luminescence levels showed significant increases in LBA-0.3MNaCl. Data are the means from three independent experiments, and error bars represent standard errors of the means. *, *P* < 0.05; **, *P* < 0.01; and ***, *P* < 0.001, ****, *P* < 0.0001 using one-way ANOVA. A PlfA-specific band was absent in the *Δplf* mutant (QT5973), whereas a strong band was detected in wild-type strain QT598 in LBA-0.3M NaCl. Positive control is PL Fimbriae in ORN172, and negative control is QT598Δ*plf*. Hemagglutination with human erythrocytes in the presence of 2.5% D-mannose confirmed that MRHA was stronger in QT598 scraped from LBA-NaCl compared to LBA, whereas the agglutination was absent in the *Δplf* mutant. Positive control is PL Fimbriae in ORN172.

**FIG. 3.**
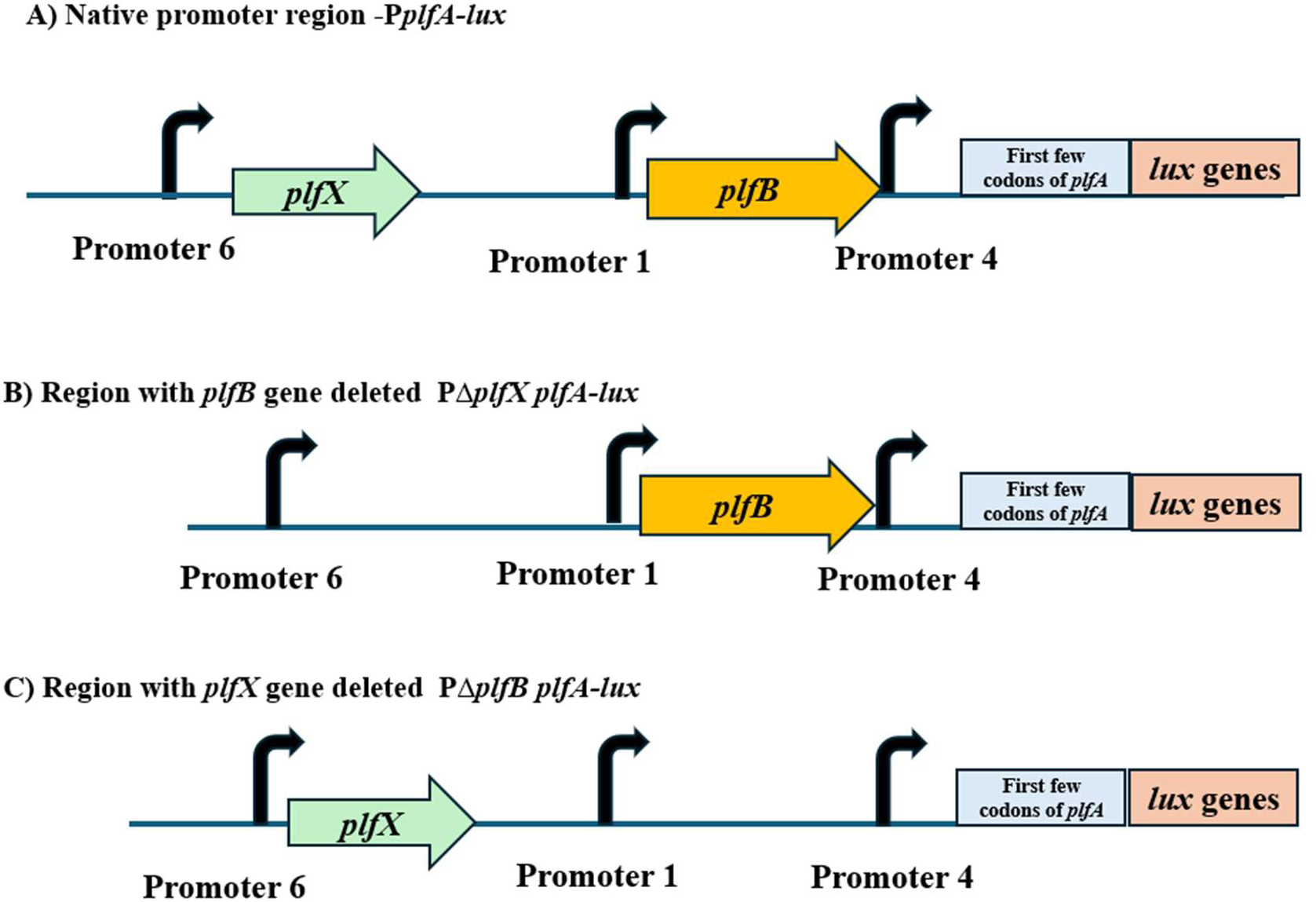
Illustration of site-specific insertion of *Pplf-lux* fusions and knock-out of predicted regulatory genes integrated at the Tn7 site of QT598. Three *E. coli* QT598 derivatives were constructed to study *plf* fimbrial regulation: (A) QT598 Pplf-lux (wild-type promoter fusion), (B) QT598 P*plf-ΔplfX-lux* (with the small *plfX* gene near the promoter removed), and (C) QT598 P*plf-ΔplfB-lux* (with the upstream *plfB* regulatory gene deleted). The schematic depicts each of the site-specific transcriptional fusions integrated at the chromosomal Tn7 site.

The decrease in the expression of *plf* in the absence of the ColV-plasmid encoded *plf* gene cluster, strain QT598Δ*plf*, showed that elements within the deleted genetic region including native regulatory region and copies of *plfB* and *plfX*, might also contribute to increased expression or positive feedback. The confirmation of production of PL fimbriae was verified by western blot using PlfA-specific antibodies (Fig. 2B). A strong band was detected in the QT598 from LBA-NaCl compared to the LBA after 3 successive passages at 37°C, which corresponds to the expected size of PlfA. Mannose-resistant hemagglutination (MRHA) with human O^+^ RBCs confirmed the high expression of PL fimbriae in salt compared to LBA in the presence of 2.5% D-mannose (Fig. 4).

**FIG. 4.**
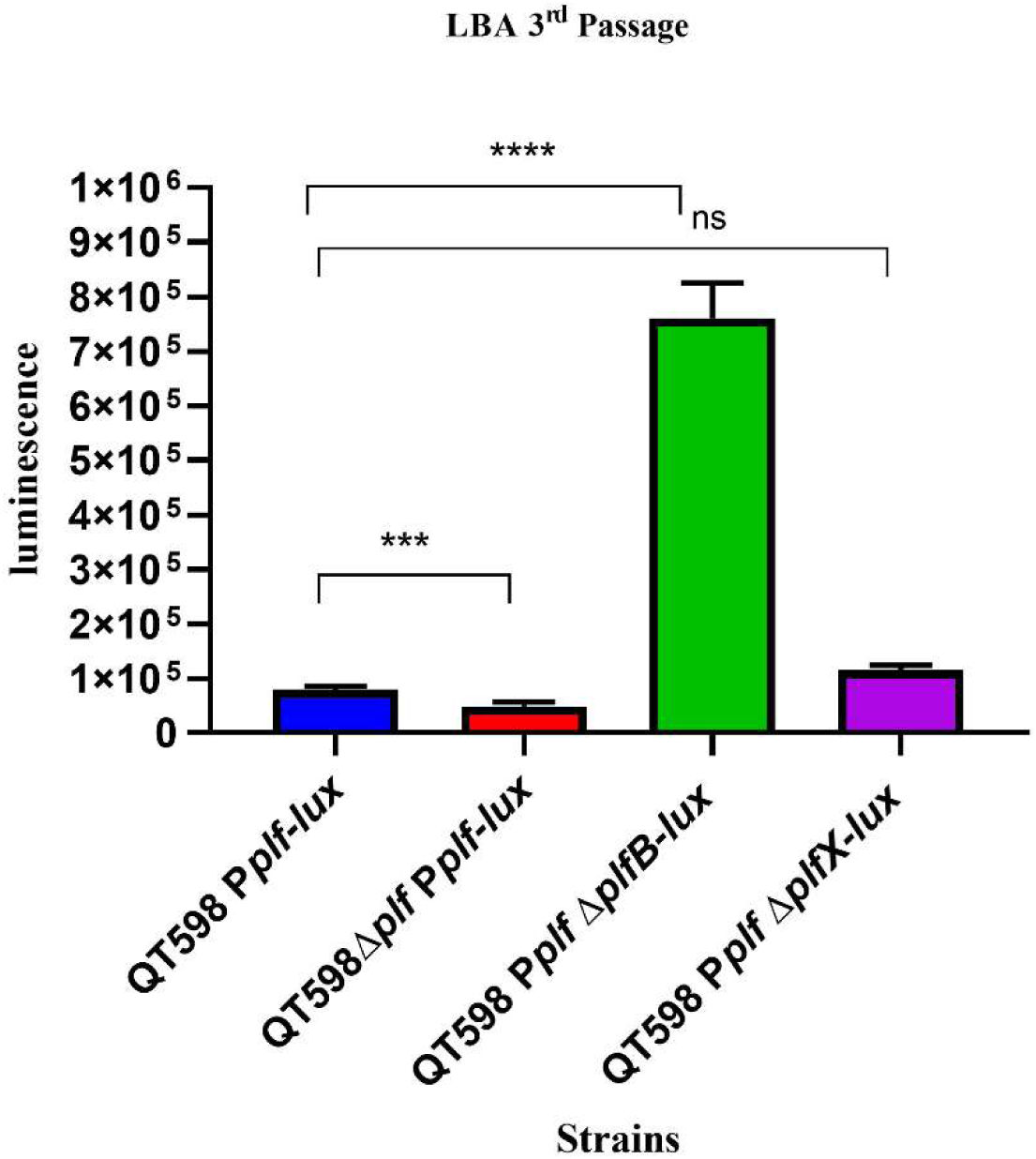
The site-specific insertion of *PplfΔplfX-lux* and *PplfΔplfB-lux* at Tn7 site of QT598. Luminescence from QT598 P*plf-lux*, QT598 P*plf**Δ**plfX-lux*, and QT598 P*plf**Δ**plfB-lux* were measured after three consecutive passages of growth on LBA. The luminescence levels showed significant increases in QT598 P*plf**Δ**plfB-lux*. Data are the means from three independent experiments, and error bars represent standard errors of the means. Ns= not significant, ***, *P* < 0.001, ****, and *P* < 0.0001 using one-way ANOVA.

### The effect of some global regulators on *plf* expression

The strains QT598 (QT5828), QT598 Δ*argR (*QT5866), QT598 Δ*hns* (QT5990), and QT598 Δ*lrp* (QT6007) were grown three times on LBA at 37°C. The luminescence analysis revealed the opposing regulatory influences of *hns* and *argR* on the *plf* promoter. The *hns* mutant had increased expression indicated H-NS is a major repressor of the *plf* promoter, based on the significant increase in the luminescence level, ∼3 times relative to QT598, which is a coordinate role of *H-NS* as a global repressor (19, 20). In contrast, the *argR* mutant showed lower luminescence than the wild type implicating ArgR as a positive regulator. This suggests that ArgR may be an accessory activator required for full promoter activity, directly or indirectly (21, 22). The *lrp* mutant demonstrated a sharp decrease in *plf* expression, which suggests Lrp likely acts as an important positive regulator (14) (Fig. 5). In line with changes in phenotypic expression, the hemagglutination was weaker for the QT598Δ*argR* and QT598Δ*lrp* mutants, but stronger for the QT598Δ*hns strain.* A PlfA -specific band was stronger in the QT598Δ*hn*s strain, whereas it was weaker in Δ*argR* strain, and there was no detectable band for the QT598Δ*plf* and QT598Δ*lrp* mutants. In silico analysis of the *plf* regulatory region identified several AT-rich sequences characteristic of potential H-NS binding sites, particularly in regions between *plfX*, and *plfB*. Further, one to two imperfect palindromic motifs similar to previously described Lrp binding sites were identified close to the promoter, suggesting that Lrp may play a direct modulatory role in *plf* expression. In contrast, no high-confidence ArgR (ARG box) motifs were identified within the analyzed sequence, making direct regulation of the *plf* operon by ArgR unlikely. The expression levels of *plfA* by strains QT598, QT598 Δ*argR,* QT598 Δ*hns*, and QT598 Δ*lrp* were also investigated by RT-PCR after 3 passages on LBA. These findings showed increased transcription in QT598 Δ*hns ∼* 6 times, whereas the other mutants demonstrated reduced expression relative to QT598 (Fig. S2).

**FIG. 5.**
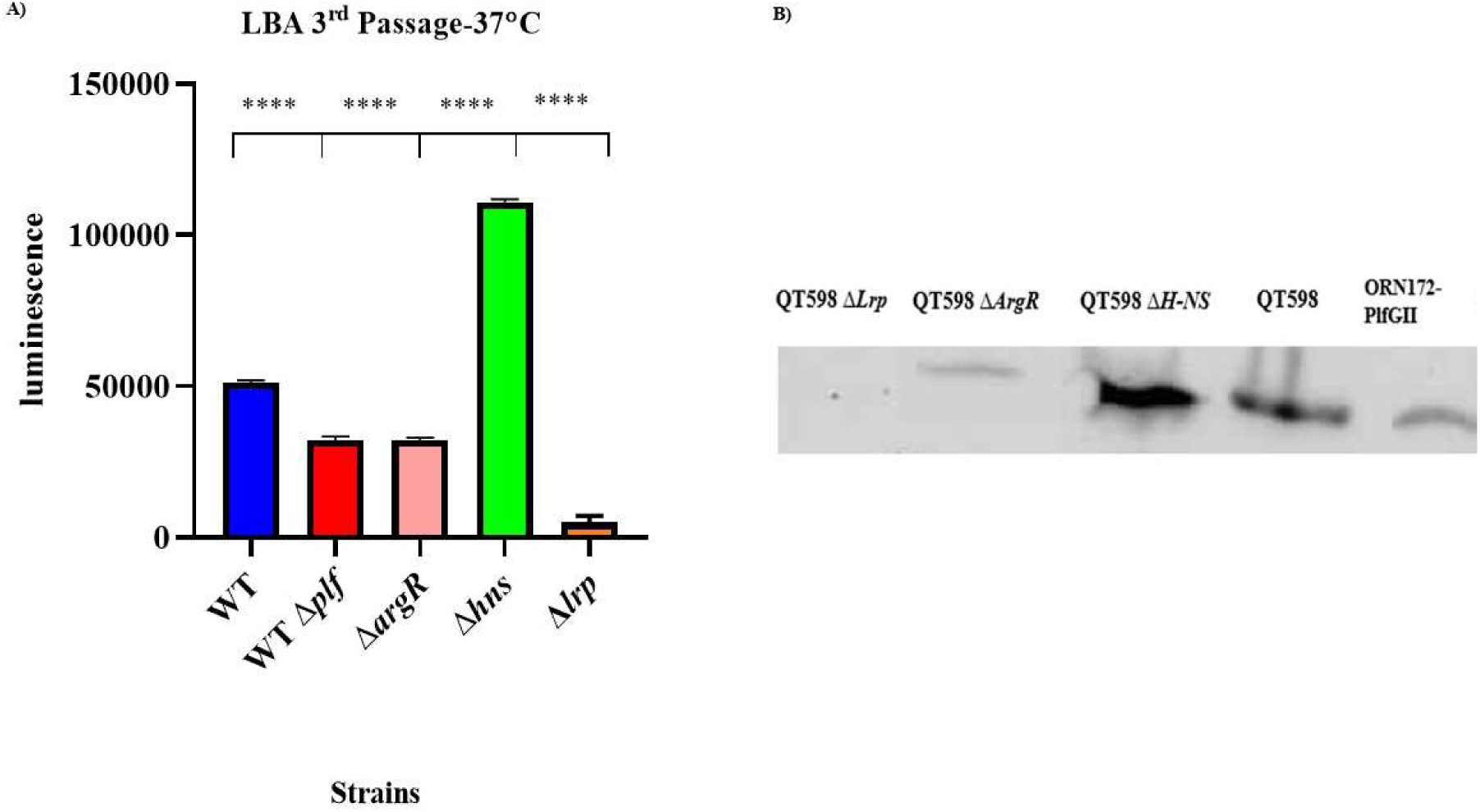
Analysis of gene-dependent regulation using *lux* reporter activity and western blot. **(A)** Luminescence from strains carrying the P*plf*-*lux* reporter was measured over three consecutive passages in LBA. Data represent mean ± SEM of biological replicates. Data are the means from three independent experiments, and error bars represent standard errors of the means. *, *P* < 0.05; **, *P* < 0.01; and ***, *P* < 0.001, ****, *P* < 0.0001 using one-way ANOVA. **(B)** Western blots specific for the PlfA protein.

### Investigation of the ratio of PL fimbriae positive cells by immunofluorescence

Bacterial fimbrial adhesins including P and type 1 fimbriae are commonly subjected to phase variation of expression, with part of a population not producing the fimbriae (OFF-phase) and some of the cells producing the fimbriae (ON-phase) (23–26). We therefore wished to investigate levels of expression or the frequency of ON-phase of cells in a specific population following growth under specific conditions. Immunofluorescence analysis was performed with strain and antibody against whole PL fimbriae including: ORN172 PlfGII (QT5726) a clone containing a plasmid with the full *plf* gene cluster from strain QT598 encoding PL fimbriae with the PlfG class II adhesin (6), strain QT598, QT598Δ*plf* (QT5973), and QT598Δ*plfB* (QT6308) following three successive passages on LBA, LB, and LBA supplemented with 0.3 M NaCl, which revealed clear differences in PL fimbrial production. For each strain and condition, three independent experiments were performed on different days, each including three technical replicates. In each replicate, 10 randomly selected microscopic fields were analyzed, with approximately 100 bacterial cells per field, yielding ∼1,000 cells per replicate and ∼9,000 cells per condition. The percentage of PL-positive cells was calculated relative to the total number of cells counted (Fig. S3).

Percent PL-positive cells were calculated as the ratio of fluorescent (PL-positive) cells to the total number of DAPI-stained bacteria in each field. The ORN172-PlfGII strain exhibited PL fimbriae in 2.45% of the bacterial population (Fig. 6A), while QT598 wild-type background demonstrated 1.12% PL-positive cells when cultivated on LBA. Notably, growth on LBA supplemented with 0.3 M NaCl increased the frequency of PL-positive cells in WT strain QT598 to 5.3% (Fig. 6B). In contrast, the Δ*plf* mutant (QT5973) exhibited only background-level signal, confirming the specificity of the antibody staining and the absence of production of PL fimbriae by this mutant (Fig.6C). Interestingly, loss of the *plfB* gene, predicted to encode a PapB-family regulator, resulted in a significant increase of PL-positive cells compared to the wild-type following growth on either LBA (4% PL-positive) or LBA 0.3 M NaCl (7% PL-positive).

**FIG. 6.**
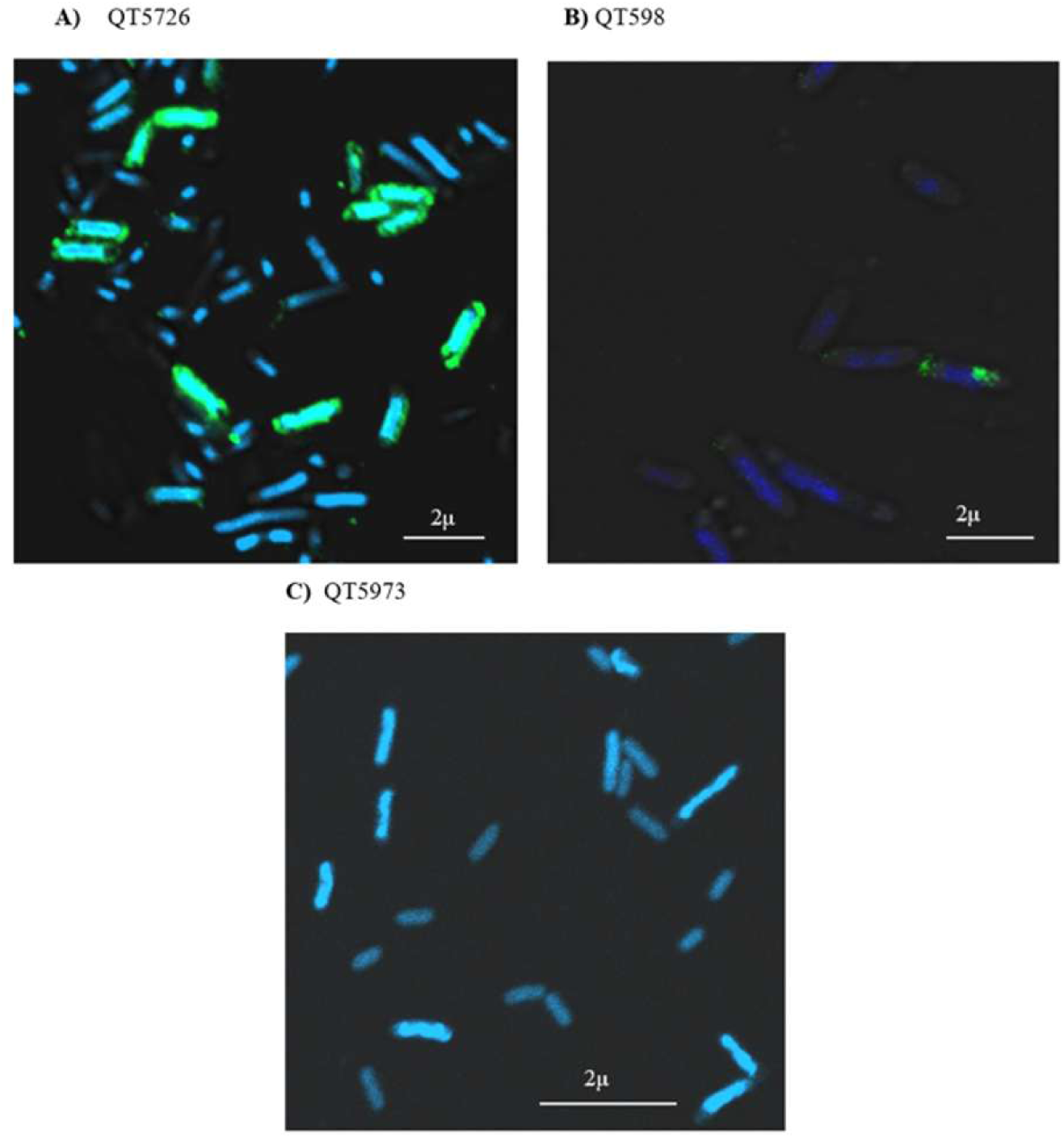
Confocal assessment of PL fimbrial ON/OFF populations. **A)** Positive control (QT5726). **B)** wildtype strain (QT598). **C)** Negative control, QT598 Δ*plf,* (QT5973). Nuclei were stained with DAPI (blue), and bacteria expressing PL fimbriae are green (Alexa Fluor 488). Images are representative of three independent experiments. Z-stack images were acquired using a Zeiss LSM 780 confocal microscope, and intensity calibration was performed using ZEN Black software.

### RT-PCR analysis of predicted promoter activity

Strain QT598 was grown on LBA for 3 passages, and total RNA was extracted and reverse transcribed. RT-PCR analysis was used to determine which of the six predicted promoter regions were transcriptionally active. Three transcriptional regions were detected, corresponding to the regions initially designated P6, P1, and P4. These regions were positioned upstream of *plfX*, *plfB*, and *plfA*, respectively, whereas no detectable transcripts were obtained for the other putative promoter regions (Fig. S4).

### Primer extension to determine the start sites of the *plf* promoters

The *plfB* gene was of particular interest because it is predicted to encode a PapB-like regulatory protein, yet its transcriptional organization was not clearly defined. Alternative annotations suggested the presence of an upstream ATG codon that could give rise to a longer *plfB* coding sequence, creating uncertainty about the correct transcriptional start site and the possible size of the predicted regulator. This uncertainty made it necessary to experimentally determine promoter usage and transcription initiation. In contrast, *plfX* is predicted to encode a small protein of unknown function adjacent to promoter 6. To directly define transcription initiation, transcriptional start sites associated with promoter 6 (*plfX*) and promoter 1 (*plfB*) were mapped using RNA extracted from strain QT598 after three successive passages on LBA. For promoter 6, a guanine residue was identified as the +1 position, with transcription initiating at TCC, whereas for promoter 1, a cytosine residue was identified as the +1 position, with transcription initiating at GAA. These transcriptional start sites are shown in Fig. 7. Together, these results confirm the activity of distinct promoters upstream of *plfX* and *plfB* and provide a clearer framework for understanding transcriptional regulation of the PL fimbrial gene cluster. Together with the RT-PCR results, these data establish distinct promoters associated with *plfX*, *plfB*, and *plfA*. Therefore, for clarity, the promoter regions initially designated P6, P1, and P4 are hereafter referred to as *PplfX*, *PplfB*, and *PplfA*, respectively.

**FIG. 7.**
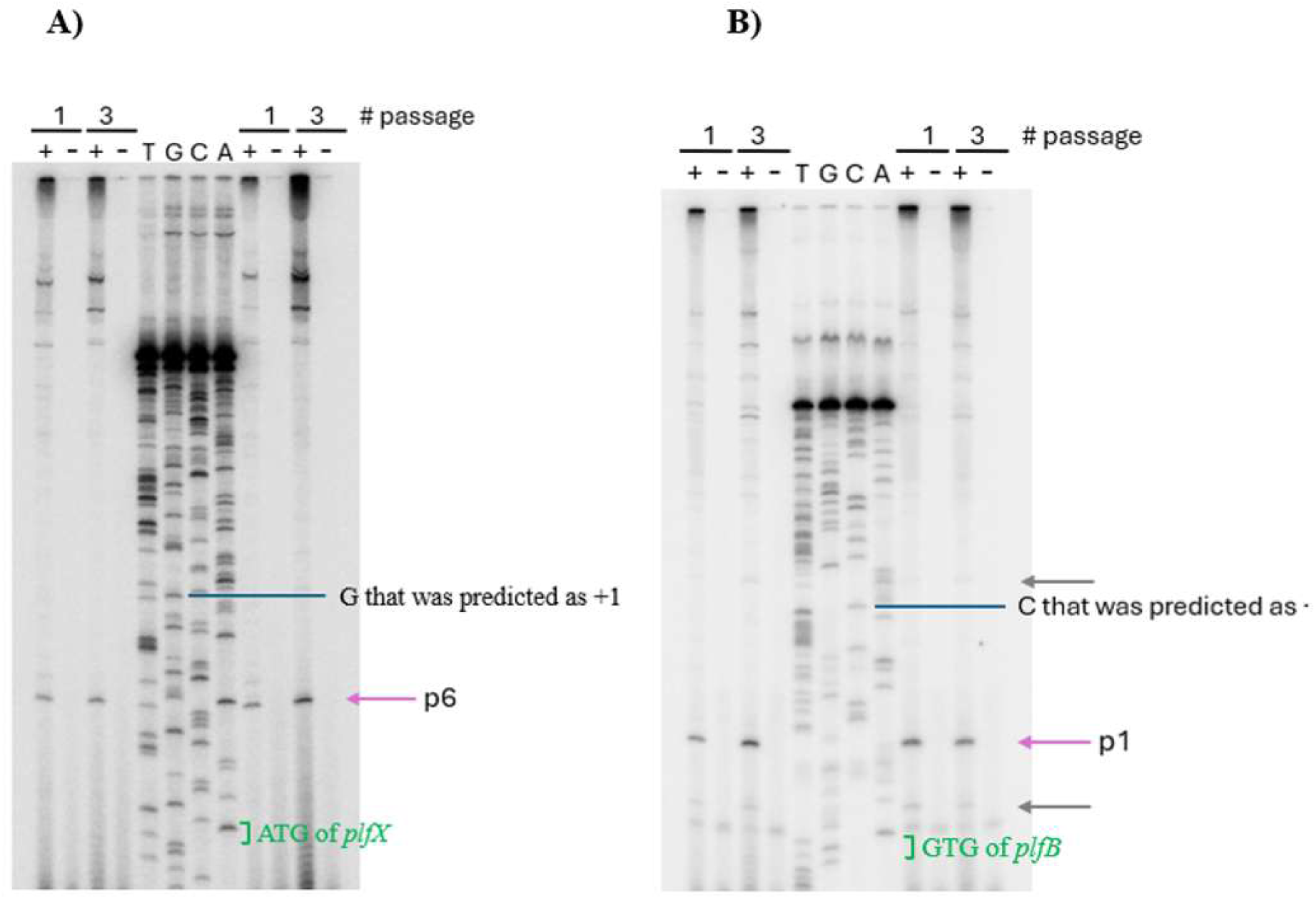
Identification of the P*plfX* and P*plfB* transcription initiation start sites by primer extension. **(A)** Foot printing to determine the *plfX* start site. **(B)** Foot printing to determine the *plfB* start site. Lanes A, C, G, and T correspond to a dideoxy sequencing reaction. Lanes + are with RT, and lanes – are without RT.

### Role of putative internal regulatory genes (*plfB* and *plfX*)

To investigate potential roles of predicted regulatory genes on Plf expression, mutants lacking either *plfX* or *plfB* genes were constructed by site-directed deletion from the pEC598 plasmid of strain QT598. These strains were grown 3 times on LBA, and the luminescence was measured (Fig. 8A). The findings showed that *plfB* acts as a repressor of *plfA* expression. Compared to the wild-type containing the P*plfA*-*lux* fusion, deletion of a single copy of *plfB* gene from the *lux* fusion caused a significant increase in luminescence. Likewise, luminescence levels in the Δ*plfB* mutant containing the complete P*plfA*-*lux* fusion (strain QT598Δ*plfB*::P*plf-lux*) were also increased. Finally, deletion of both the native *plfB* and *plfB* gene within the *lux* fusion promoter resulted in highest levels of luminescence that were 3-fold higher than the *lux* fusion strain containing 2 copies of *plfB* (Fig. 8A). In contrast to the loss of *plfB*, loss of either one or both copies of the *plfX* gene in the *lux* fusion strains resulted in a significant reduction in the levels of luminescence with a reduction to ≈50% in the strain lacking both copies of *plfX* (Fig. 8B). To confirm a role for these potential *plf* regulatory genes, derivatives of QT598 lacking native copies of either *plfX* or *plfB* were grown for 3 consecutive passages on LBA, and PL fimbriae were extracted for verification of production levels by western blot. The *plfB* mutant was found to produce more PL fimbriae, and its hemagglutination activity was higher than that of the wild-type strain, which is consistent with the *lux* reporter experiments (Fig. 8B).

**FIG. 8.**
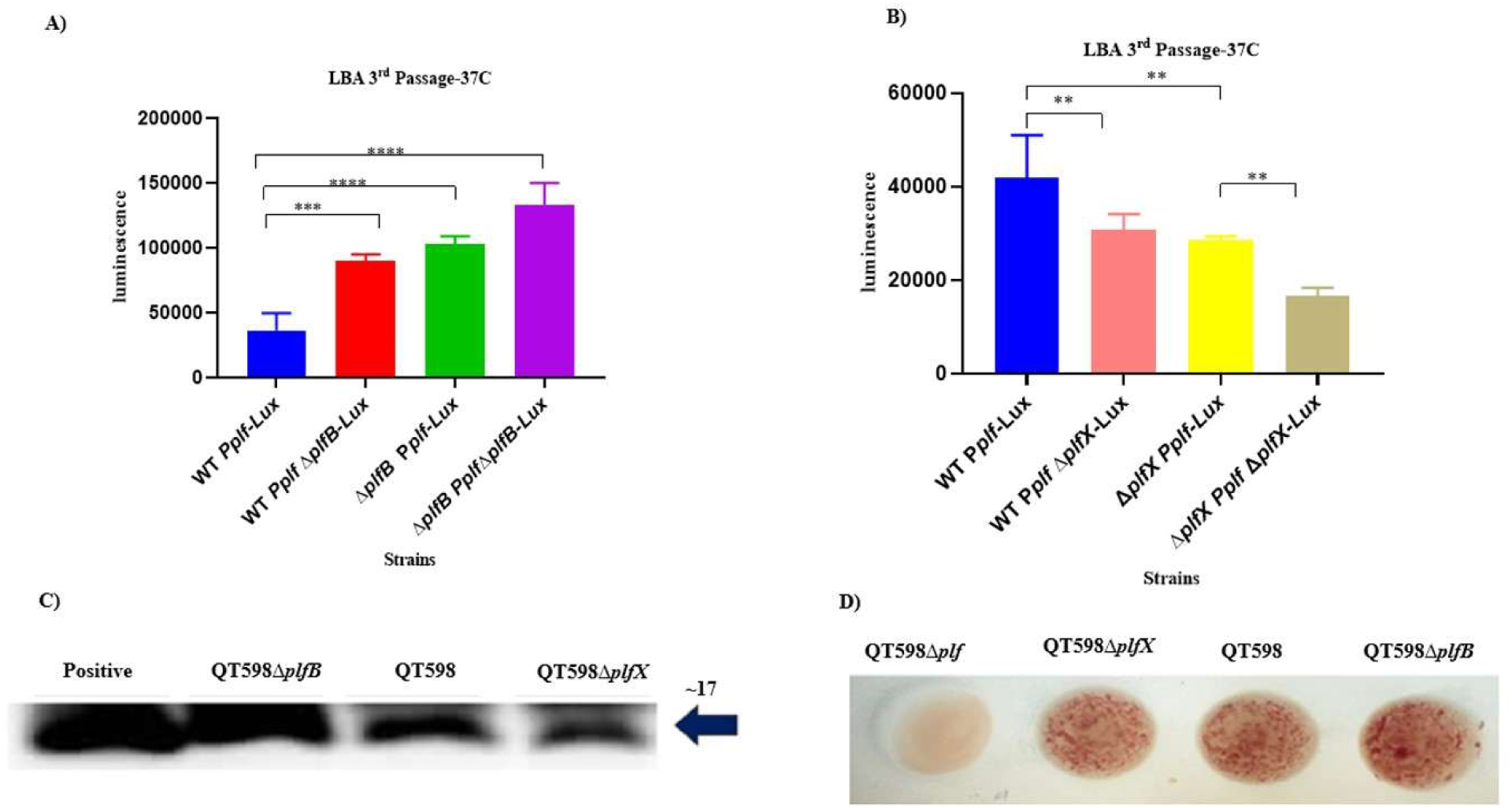
Role of internal regulators (PlfB and PlfX) in PL fimbriae expression. **A)** Comparative luminescence levels in strains containing two, one, or no copies of either *plfB* **(A)** or *plfX* **(B)** genes after three passagers on LBA (mean ± SEM, biological replicates). Data are the means from three independent experiments, and error bars represent standard errors of the means. *, *P* < 0.05; **, *P* < 0.01; and ***, *P* < 0.001, ****, *P* < 0.0001 using one-way ANOVA. **(C)** Western blot for PlfA protein demonstrated greater levels of PL fimbriae for the Δ*plfB* mutant, whereas PL fimbriae production for the Δ*plfX* mutant was similar to wild-type strain QT598. **(D)** Macro-hemagglutination of human O⁺ erythrocytes with 2.5% D-mannose. There was no difference in hemagglutination between QT598 and QT598Δ*plfX,* whereas the level of agglutination by the Δ*plfB* mutant was stronger.

### Loss of the *plfB* regulatory gene increases lung colonization during turkey infection

Since loss of the *plfB* regulator demonstrated an increased level of production of PL fimbriae, we were curious to investigate if this regulator could influence colonization in a turkey respiratory infection model. We hypothesized that PL fimbriae-and their regulation by *plfB*-may serve a host-adapted role in tissue colonization. To assess the role of *plfB* during APEC infection in turkeys, we generated a *plfB* deletion mutant and a complemented strain in which the *plfB* gene was reintroduced into the chromosome. The verification of changes in Plf were verified by Western blotting using PlfA-specific antibodies (Fig. S5) and mannose-resistant hemagglutination (MRHA) assays. The PlfA subunit was detected in wild-type and complemented strains at similar levels but was increased in the Δ*plfB* mutant, demonstrating successful complementation. Similarly, MRHA was restored in the complemented strain but reduced in the Δ*plfB* mutant, confirming functional recovery of the *plfB* gene. We then tested whether the regulatory effect of *plfB* observed *in vitro* had any appreciable effect on infection in the natural host model.

At 48 h post-infection, the wild-type strain QT598 colonized the lungs of turkey at levels between ∼10^6^ to 10^7^ CFU/g. Interestingly, however, the Δ*plfB* mutant was recovered from the lungs of infected poults with colony-forming units (CFU) at significantly higher numbers (P < 0.05) ranging between ∼10^8^ to 10^9^ CFU/g (Fig. 9A). The complementation of the Δ*plfB* mutant with *plfB* in single-copy on the chromosome (QT6335) restored colonization levels in the lung to levels comparable to the wild-type strain. These results indicate that deletion of *plfB* leads to a significant increase in colonization of turkey lungs, which may potentially be due to greater levels of expression of PL fimbriae in lung tissues. Bacterial burdens in the liver and spleen were also determined 48 h post-infection and were similar for all strains with a mean ≈ 10^5^ CFU/g in the livers (Fig. 7B), and ≈ 10^4^ CFU/g in the spleens (Fig. 9C). These data indicate that loss of the *plfB* gene had no appreciable effect on colonization of these systemic organs compared to colonization of the lungs. Despite high bacterial numbers in multiple organs, none of the turkeys exhibited typical clinical signs of colibacillosis, such as respiratory distress, lethargy, or huddling. Blood samples were also tested for bacterial numbers, but no bacteremia was observed in any of the groups. Overall, results suggest that *plfB* may potentially decrease levels of expression of PL fimbriae in lungs and that this may be linked to a reduction of colonization levels in the lung tissues of turkeys compared to the strain lacking *plfB*.

**FIG. 9.**
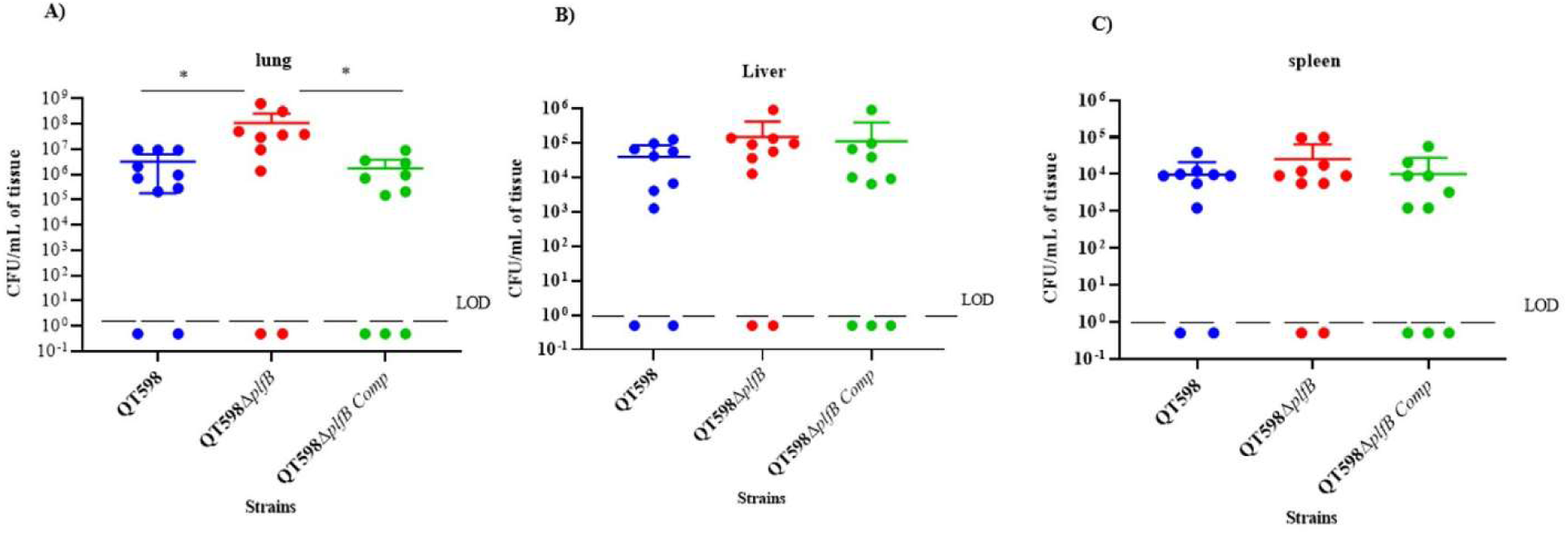
Bacterial colonization of turkey tissues following infection with wild-type QT598, Δ*plfB* mutant, and complemented strains. Six-day-old turkey poults were inoculated, and viable bacterial counts were determined from the right lungs (A), livers (B), and spleens (C) at 48 h post-infection. Each point represents an individual tissue sample from a poult, with horizontal lines indicating median CFU/g. The limit of detection (LOD) is shown by a dashed line. In the lungs, deletion of *plfB* resulted in a significant increase in colonization relative to the wild-type strain (*, P < 0.05). No significant differences (ns) were detected among strains in the liver or spleen. *, P < 0.05; **, P < 0.01; ***, P < 0.001 (one-way ANOVA).

### *In vivo* regulation of PL fimbrial expression by *plfB* during turkey lung infection

Based on the increased bacterial burdens observed in the lungs of turkeys infected with the Δ*plfB* mutant, we next examined whether this increase was accompanied by altered expression of plfA, which encodes the major PL fimbrial subunit. To do so, *plfA* transcript levels were quantified by qRT-PCR in lung tissues from poults infected with either the wild-type strain QT598 or the QT598 Δ*plfB* mutant. Importantly, expression levels were normalized to the housekeeping gene *rpoD* to control for differences in bacterial load between samples. This normalization ensures that changes in *plfA* transcript abundance reflect transcriptional regulation rather than simply increased bacterial numbers in the lungs. Five individual lung samples from each group (n = 5 per strain) were analyzed independently.

Following normalization, *plfA* expression was approximately 10-fold higher in lungs infected with the Δ*plfB* mutant compared to the wild-type strain (Fig. 10). These results support a role for PlfB as a negative regulator of PL fimbrial expression during respiratory infection. This result suggests that PlfB could help to modulate levels of PL fimbrial expression in lung tissues, which contain host cell receptors recognized by the PlfG adhesin (18). The lung tissues and environmental conditions such as temperature, osmolarity, host factors, or nutrient availability may naturally promote *plfA* expression, which, however, may still be repressed by PlfB. As such, the removal of the *plfB* regulatory gene still resulted in a further increase in *plf* expression and enhanced colonisation in the turkey lungs. We have previously determined that comparative expression of *plfA* in the lungs of infected turkeys was higher than its expression *in vivo* after three consecutive passages on LB agar (18). As such, we were interested in verifying if *plfB* transcript levels were downregulated in the lungs compared to growth in vitro. We compared *plfB* expression after 3 successive passages on LBA, in comparison to *plfB* expression from infected turkey lung. Quantitative RT-PCR analysis, normalized to the housekeeping gene *rpoD*, revealed a clear difference in *plfB* expression between infected lung samples and bacteria grown on LBA. In lung tissues, *plfB* expression was consistently low, with ΔCt values clustering close to zero (range: 0.58–1.18; mean ΔCt ≈ 0.89), indicating limited transcription under these conditions. In contrast, growth on LBA resulted in markedly lower ΔCt values (range: −9.07 to −10.97; mean ΔCt ≈ −9.83), reflecting a strong induction of *plfB* expression *in vitro* (Fig. S6).

**FIG. 10.**
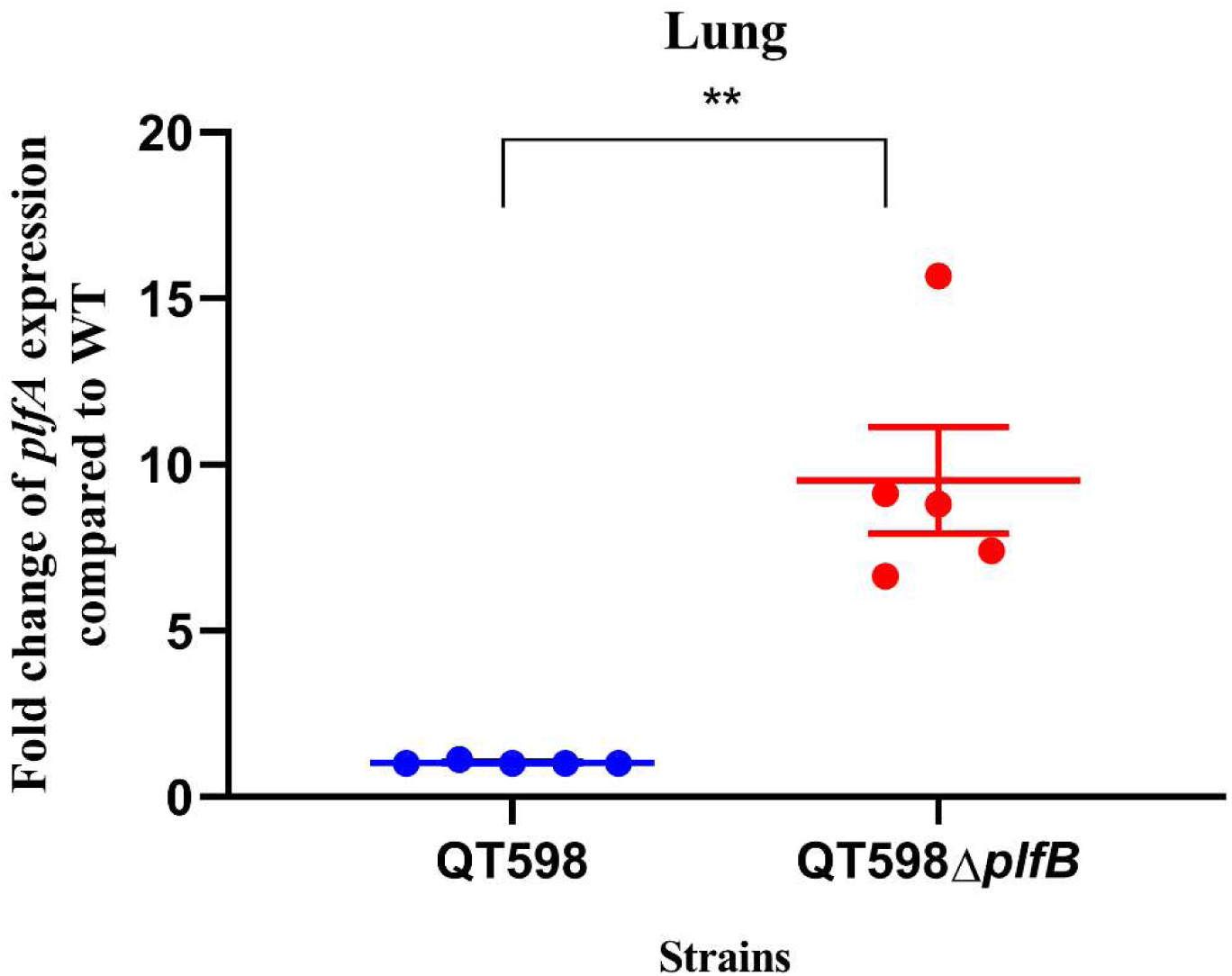
The expression of *plfA* in the lung of infected turkey poults with QT598 and QT598Δ*plfB*. Quantitative RT-PCR analysis showing the fold change in *plfA* expression in strains recovered from lung tissues. Expression of *plfA* was significantly higher in the QT598**Δ***plfB* compared to the QT598. Data represent the mean ± standard errors of the mean (SEM) from 3 biological replicates.

When considered together with our previous analysis of *plfA* expression, these data reveal an inverse relationship between *plfB* and *plfA* expression across growth conditions. In the lung environment, a *plfB* mutant had *increased plfA* transcript levels, consistent with increased production of PL fimbriae during infection. Conversely, during growth on LBA, *plfB* expression was strongly induced, while *plfA* expression was comparatively lower. Despite observing increases in the levels of PL fimbriae due to loss of *plfB*, other regulatory cues are likely to play an important role in differences of expression *in vivo* compared to in vitro. However, based on results obtained from expression of PL fimbriae in the *plfB* mutant during in vitro growth or on the increases in *plfA* expression levels observed in tur key lungs during infection, results PlfB seems to play a role as a negative regulator of PL fimbriae both *in vitro* and *in vivo*.

### Electrophoretic mobility shift assays identify *PlfB* as a promoter-binding regulator

To determine whether PlfB and PlfX directly interact with the regulatory region of the *plf* gene cluster, we performed EMSA using purified PlfB or PlfX proteins and a 6-carboxyfluorescein (56 FAM) labelled fragment encompassing the *plf* promoter regions. Each protein was extracted and purified. The *lacI* promoter fragment was included as a negative control and incubated with proteins. Either PlfB or PlfX was incubated with 3 different DNA fragments including the *PplfX* promoter region (promoter 6), the *PplfB* promoter region (promoter 1), and *PplfA* promoter region (promoter 4) (Fig. S7). EMSA results showed that PlfB bound specifically to the *PplfA* promoter fragment, while no binding to either the *PplfB* or *PplfX* promoter fragments was observed. Further, neither of these proteins bound to the *lacI* promoter region that was used as a negative control (Figs. 10 and 11). In contrast to PlfB, PlfX did not show detectable binding to any of the tested DNA fragments under these conditions (Fig. 11). These findings indicate that PlfB functions as a direct DNA-binding protein targeting the *plfA* promoter, consistent with its role as a repressor of the *plf* gene cluster. In contrast, the lack of detectable DNA binding by PlfX suggests that its regulatory function may not involve direct interaction with the different *plf* gene promoter regions, or that it requires additional cofactors or specific conditions not present in our assay.

**FIG. 11.**
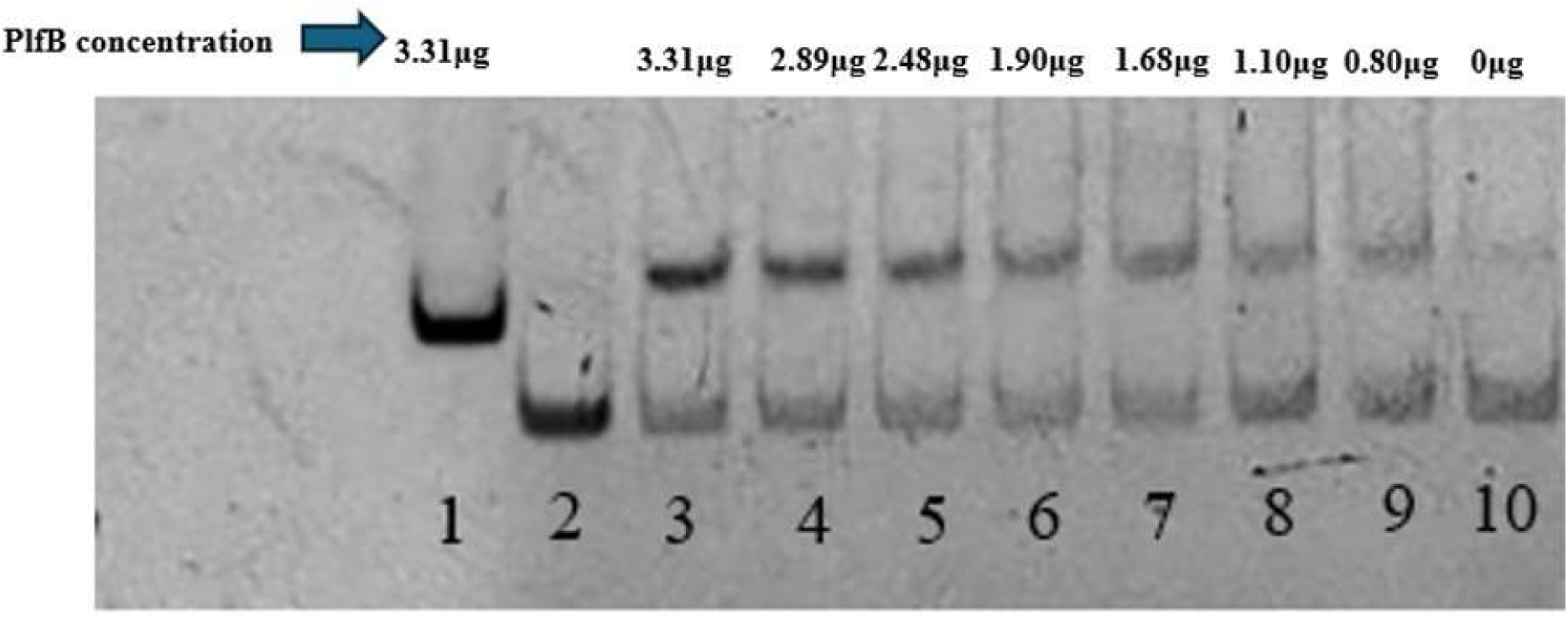
Electrophoretic mobility shift assay (EMSA). EMSA was performed using varying concentrations of PlfB protein incubated with 30 ng of the *plfA* promoter region. Lane 1: *LacI* promoter (300 bp) as a negative control with plfB protein; Lane 2: *plfA* promoter (200 bp) alone. Lanes 3–10: PlfB– *plfA* promoter interactions at decreasing protein concentrations. The upper bands correspond to the PlfB– DNA complexes, while the lower bands represent free DNA. No binding was observed when either the *plfB* or *plfX* promoter regions were used.

**FIG. 12.**
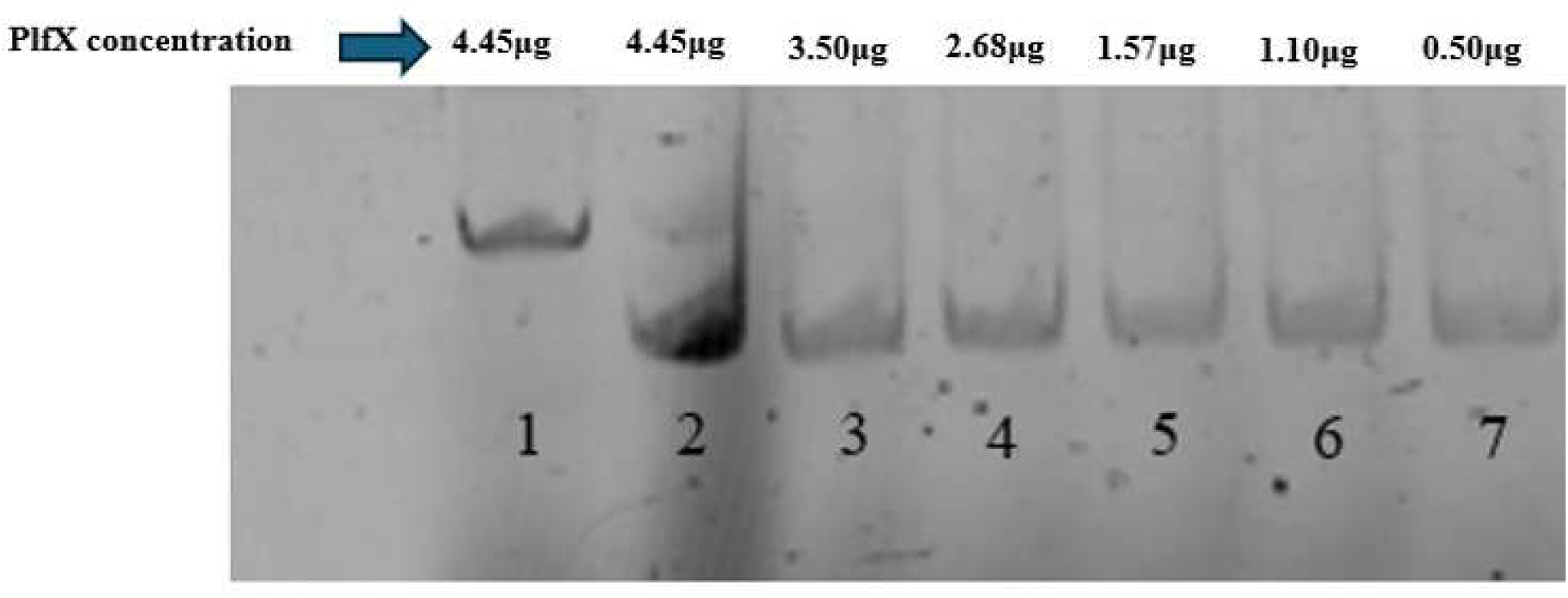
Electrophoretic mobility shift assay (EMSA). EMSA was performed using varying concentrations of PlfX protein incubated with 30 ng of the *plfA* promoter region. Lane 1: *LacI* promoter (300 bp) as a negative control; Lane 2: *plfX* promoter (200 bp) alone. Lanes 3–7: represent free DNA. There was no shift. Similarly, no binding was observed when either the *plfB* or *plfX* promoter regions were used.

## Discussion

PL fimbriae are a recently described group of fimbrial adhesins belonging to the π fimbrial family, which have been identified in the genomes of some *E. coli* strains associated mainly with extra-intestinal infections in humans and poultry (6, 18). *E. coli* strain QT598 isolated from a young turkey poult, encodes PL fimbriae containing a PlfG class II adhesin that recognizes α1-2 glycans such as Lewis b, Lewis y, and H antigens, which were shown to promote adherence to human uroepithelial cells and turkey lung tissues, and contributed to respiratory infection and colonization of turkey lungs in an experimental infection model. In strain QT598, the PL fimbriae are encoded by the *<u>plf</u>* genes located on a ColV-like plasmid called pEC598, however their expression was found to be limited *in vitro* and required multiple passage on LB agar plates to increase production of these adhesins. Despite limited expression in vitro, *plf* gene expression was increased during infection in both murine urinary tract and turkey respiratory tract models and contributed to infection. As little information is available concerning conditions influencing PL fimbriae production or regulation, the purpose of this study was to investigate environmental and regulatory aspects influencing PL fimbriae production and gene expression.

To investigate, the effect of different cultural conditions of *plf* gene expression, a single-copy *lux* operon was fused to the *plf* promoter region encompassing a 1550 bp region that was found to contain 3 functional promoters (Fig. 3A) specific for each of the two putative regulatory genes, *plfB* and *plfX,* encoding a paralog of the PapB regulator and a gene of unknown function, respectively. Further, this region includes the intergenic and promoter region upstream of the *plfA* promoter, which likely contains multiple binding-sites for global and internal regulators of *plf* gene expression. By using this system, we were able to investigate levels of *plf* expression following growth under different conditions. Results indicated that *plf* expression was increased at higher temperatures (37° and 42°C) compared to 25° or 30°C, and LBA at 37° C was shown to increase *plf* expression compared to other media tested. Interestingly, increasing NaCl in the medium to 0.3 M resulted in a marked increase in *plf* expression (Fig. S1) as well as PL fimbrial production (Fig. 2). Interestingly, increased NaCl has also been shown to decrease expression of some fimbriae such as type 1 fimbriae and curli, while increasing levels of expression of other fimbriae, such as P and F1C fimbriae in UPEC strain CFT073 (27, 28). Thus, Plf expression may similarly be favored under higher salt conditions, such as in the urinary tract during infection (6). However, other conditions or host environmental cues are also likely to upregulate Plf within avian respiratory tissues, and this has similarly been shown to be the case for P fimbriae in APEC (29).

In addition to local or internal regulators, such as PapB and PapI or their paralogs, that are linked to specific fimbrial gene clusters such as *pap*-and *foc*-encoded fimbriae, the regulatory control of fimbrial systems that are dependent on PapB family regulators comprise epigenetic mechanisms that include global regulatory proteins such as H-NS, Lrp, CRP, and changes in GATC methylation sites that are targets for the Dam methylase protein (30) and can repress both type 1 and P fimbriae, particularly at lower temperatures which favor H-NS binding to DNA target sequences within promoter regions (11, 16, 17, 20). By contrast, leucine responsive regulatory protein (Lrp) which plays a key role in phase-variation switching, was shown to be necessary for optimal expression of multiple types of fimbriae including P, type 1, and F1C fimbriae (11, 17, 30). Its role has been shown to be dependent on nutrient status and amino acid availability as a global regulator and DNA-binding interactions that can block Dam methylation to promote expression of different fimbrial systems. As such, it is not surprising that a *hns* mutant of strain QT598 demonstrated increased levels of plf expression and PL fimbriae production, whereas a *lrp* mutant demonstrated a strong decrease in *plf* expression (Fig. 5). Although specific mechanisms of how these global regulators control *plf* expression have not been determined, it will be of interest to further elucidate their specific roles as well as the possible importance of other regulators such as CRP and Dam methylase in controlling *plf* gene regulation.

The local regulatory control of PL fimbriae is distinct compared to most other systems that include a PapB family paralog, since most of these systems also encode a PapI-related protein. In the case of the Pap system, these two proteins play complementary roles in fimbrial regulation with PapB playing a role in expression of the PapI protein, which also positively regulates PapB and expression of the fimbrial structural genes (16, 30, 31). Since the *plf* system lacks a PapI equivalent, but only encodes a PapB paralog, its regulation is atypical compared to some other fimbrial systems regulated by PapB-like regulators. Further, unlike most PapB paralogs, loss of *plfB* was found to increase expression of *plf* and production of PL fimbriae by nearly 4-fold (Fig. 8). This contrasts with most other fimbrial systems under the regulatory influence of a PapB paralog, wherein loss of the regulatory protein eliminates or strongly reduces fimbrial expression. For either of the Pap, F1C, or Sfa fimbriae, which have highly similar regulatory proteins, loss of PapB, FocB, or SfaB respectively results in a strong decrease in fimbrial expression (32–34). Similarly, in *Salmonella enterica* PefB acts as a positive regulator to promote Pef expression (5). In the case of Afa-3 adhesin, which lacks a PapI-like regulator, loss the AfaA regulator by disruption of the *afaA* gene eliminated phenotypic expression of Afa adhesin and loss of hemagglutination (35).

For K99 fimbriae, which has two PapB family proteins, FanA and FanB, both regulators were important for optimal expression as frameshift mutations in either of these genes reduced expression levels by at least 10-fold (36). Among the PapB paralogs, the only protein shown to exert a clear negative effect on fimbrial expression was ClpB of CS31A fimbriae (37). Loss of *clpB* resulted in increased expression of the CS31A-encoding genes, and ClpB was shown to act as a co-repressor stabilising Lrp-mediated repression (37). As PlfB was shown to bind directly to the *plfA* promoter region (Fig. 9), it likely acts directly as a negative regulator of PL fimbriae. It will therefore be of further interest to determine what other regulators may influence expression of either *plfB* or *PlfA* genes and influence plf fimbrial exression in future studies.

Comparison of *plfB* expression *in vitro* and *in vivo* revealed higher transcript levels on LBA and significantly reduced expression in lung tissue. This decrease correlates with the enhanced expression of *plfA* in the lung and is consistent with a model in which host environmental conditions relieve PlfB-mediated repression, enabling elevated fimbrial production during host colonization. Such modulation reflects the tissue-specific nature of fimbrial regulation in APEC and parallels regulatory strategies observed in other pathogens adapting to host niches. The fact that there was still a higher level of *plfA* expression in turkey lungs (Fig. 10), however, indicates that the PlfB regulator may still be sufficiently present to exert some decrease in the level of *plf* gene expression and that this may have influenced levels of colonisation in the turkey lungs. Since some PapB family regulators have also been shown to affect regulation of heterogeneous fimbrial systems through crosstalk mechanisms. For example, in uropathogenic *E. coli*, overexpression of *PapB* inhibited recombination-mediated switching of the type 1 fimbrial operon’s fim switch, thereby preventing simultaneous expression of multiple fimbrial types in a single cell (3, 38, 39). It will therefore be important to more extensively verify if deletion or inactivation of some or all the elements in the *plf* system may influence the expression of other fimbrial adhesins under different growth conditions or during infection or colonisation of the host.

This study provides the first investigation of transcriptional regulation, environmental responsiveness, and in vivo role of the PlfB regulator for the Plf system of APEC QT598. Through an integrated approach combining promoter mapping, reporter assays, mutational analyses, qRT-PCR, EMSA, and a turkey infection model, we have demonstrated both global and local regulators contributing to some aspects of *plf* gene expression. Importantly, in contrast to Pap and similar fimbrial systems regulated by PapB paralogs, PlfB was shown to play a distinct role as a negative regulator of plf expression that demonstrated direct binding to the *plfA* promoter region. The archetypal PapB binds to AT-rich upstream activating regions in the *pap* promoter, modulating transcription and phase variation (31, 40). Homologs such as FocB and SfaB share similar helix-turn-helix structures and maintain conserved regulatory functions across fimbrial operons (3, 32). Structural analyses of FocB further highlight the conserved DNA-binding architecture within this family (32). Consistent with these features, we showed that PlfB binds directly to the *plfA* promoter which results in repression of transcription and decreased PL fimbrial production.

Overall, our work identifies *plfB* as a PapB-family transcriptional regulator that modulates PL fimbrial expression through direct promoter binding, environmental responsiveness, and that interplay with global regulators, such as H-NS and Lrp, contribute to Plf regulation. Comparative analysis with classical PapB-family systems suggests that while PlfB retains conserved regulatory mechanisms, the *plf* gene regulation may have diverged to meet the demands of avian colonization, potentially optimized for respiratory colonization rather than systemic infection. However, since the Plf system is also able to mediate adherence to human uroepithelial cells and is associated with *E. coli* strains isolated from urinary tract infections, it may also play a role in UTI pathogenesis for some strains (6, 18). This study also further highlights the complexity of fimbrial regulation in pathogenic *E. coli*. It establishes a foundation for further investigation of regulatory crosstalk and host-specific signals that may drive niche adaptation. This adaptive capacity of expressing multiple types of fimbrial adhesins likely contributes to the ecological success of pathogenic *E. coli* in both avian and mammalian hosts and underscores the evolutionary adaptability of fimbrial regulation within the *Enterobacteriaceae*.

## Materials and Methods

Table 1 presents the bacterial strains and plasmids used in this study. APEC strain QT598 and its derivative strains were grown at 37°C on solid or liquid Luria-Broth (LB) medium (Alpha Bioscience, Baltimore, MD, USA). The final concentrations of 100 µg/ml ampicillin, 30 µg/ml chloramphenicol, 100 µg/mL carbenicillin, and 50 µg/ml kanamycin were used when required. For the turkey infection, brain heart infusion (BHI) broth (Alpha Bioscience, Baltimore, MD, USA) was used for the growth of QT598 and its derivatives.

**TABLE 1.** List of strains and plasmids.

| Strain, plasmid, or clone | Characteristic(s) | Reference |
| --- | --- | --- |
| **Strains** |  |  |
| *E. coli* MGN-617 | *thi thr leu tonA lacY glnV supE ΔasdA4 recA:: RP42-Tc::Mu [pir]; Km<sup>r</sup>::RP4* | (6, 50) |
| ORN172 | *fim*-negative strain; *thr-1 leuB thi-1Δ(argF-lac)U169 xyl-7 ara-13 mtl-2 gal-6 rpsL tonA2supE44Δ(fimBEACDFGH)::Km pilGI* | (6) |
| QT598 | APEC O1: K1 (Sequence type, ST1385) | (6, 50) |
| QT2087 | MGN-617 + pLOF / Km; Amp<sup>r</sup> Km<sup>r</sup> | (50) |
| QT5726 | ORN172/pIJ497 (reference clone PL fimbriae with PlfGII adhesin) | (6) |
| QT5732 | ORN172/ pUCmT empty vector | (6) |
| QT5765 | QT598(pSTNSK) Tp<sup>r</sup> | This study |
| QT5792 | QT598 $\Delta argR::$ kan, Km<sup>r</sup> | This study |
| QT5794 | QT5792 (pSTNSK) Km<sup>r</sup> | This study |
| QT5814 | χ7213+pGP-Tn7-Cm-*Pplf* promoter region *luxCDABE*, Ap<sup>r</sup>, Cm<sup>r</sup>, Km<sup>r</sup> | This study |
| QT5828 | QT598::Tn7T-Cm::*Pplf* promoter *luxCDABE*, Cm<sup>r</sup> | This study |
| QT5866 | QT598 $\Delta argR::$ Tn7T-Cm::*Pplf* promoter *luxCDABE*, Cm<sup>r</sup> | This study |
| QT5918 | QT598 $\Delta hns::$ kan, Km<sup>r</sup> | This study |
| QT5951 | QT598 (pSTNSK) Km<sup>r</sup> | This study |
| QT5973 | QT598 $\Delta plf::$kan, Km<sup>r</sup> | This study |
| QT5990 | QT598$\Delta hns::$ Tn7T-Cm::*Pplf* promoter *luxCDABE*, Cm<sup>r</sup> | This study |
| QT5993 | QT5973 (pSTNSK) T<sub>p</sub><sup>r</sup> | This study |
| QT5995 | QT598 $\Delta plf::Tn7T$-Cm::P<sub>plf</sub> promoter *luxCDABE*,<br>Cm<sup>r</sup> | This study |
| QT5998 | QT598 $\Delta lrp::kan$, Km<sup>r</sup> | This study |
| QT6007 | QT5998 (pSTNSK) T<sub>p</sub><sup>r</sup> | This study |
| QT6016 | QT598 $\Delta lrp::Tn7T$-Cm::P<sub>plf</sub> promoter *luxCDABE*,<br>Cm<sup>r</sup> | This study |
| QT6040 | MGN-617 (pIJ621); Gm<sup>r</sup> | This study |
| QT6223 | $\chi7213$+Tn7T-Cm:: P<sub>plf</sub>$\Delta plfB$ promoter *luxCDABE*,<br>Cm<sup>r</sup>, DAP- | This study |
| QT6245 | $\chi7213$+Tn7T-Cm:: P<sub>plf</sub>$\Delta plfX$ promoter *luxCDABE*,<br>Cm<sup>r</sup>, DAP- | This study |
| QT6269 | BL21 (DE3):: PlfB, Km<sup>r</sup> | This study |
| QT6277 | $\chi7213$+Tn7T-Cm:: P<sub>plf</sub>$\Delta plfX$ $\Delta plfB$ promoter<br>*luxCDABE*, Cm<sup>r</sup>, DAP- | This study |
| QT6303 | QT598 conjugation with pMEG375 *orfE*-Gm-P<sub>plf</sub><br>$\Delta plfX$ | This study |
| QT6304 | QT598 conjugation with pMEG *orfE*-Gm-P<sub>plf</sub> $\Delta plfB$ | This study |
| QT6308 | QT598 $\Delta plfB$ | This study |
| QT6309 | BL21 (DE3):: PlfX, Km<sup>r</sup> | This study |
| QT6311 | QT6308 (pSTNSK) Km<sup>r</sup> | This study |
| QT6314 | QT598 $\Delta plfB::Tn7T$-Cm::P<sub>plf</sub> promoter *luxCDABE*,<br>Cm<sup>r</sup> | This study |
| QT6315 | QT598 $\Delta plfB$:: Tn7T-Cm:: *Pplf*$\Delta plfX$ promoter<br>*luxCDABE*, Cm<sup>r</sup> | This study |
| QT6316 | QT598 $\Delta plfB$:: Tn7T-Cm:: *Pplf*$\Delta plfB$ promoter<br>*luxCDABE*, Cm<sup>r</sup> | This study |
| QT6318 | χ7213+Tn7T-Cm:: *plfB*, Cm<sup>r</sup>, DAP- | This study |
| QT6319 | χ7213+Tn7T-Cm:: *plfX*, Cm<sup>r</sup>, DAP- | This study |
| QT6335 | QT598 $\Delta plfB$:: *plfB* Gm<sup>r</sup> | This study |
| QT6349 | QT598 $\Delta plfX$ | This study |
| QT6350 | QT6349 (pSTNSK) Km<sup>r</sup> | This study |
| QT6355 | QT598 $\Delta plfX$:: Tn7T-Cm:: *Pplf* promoter<br>*luxCDABE*, Cm<sup>r</sup> | This study |
| QT6356 | QT598 $\Delta plfX$::Tn7T-Cm:: *Pplf*$\Delta plfX$ promoter<br>*luxCDABE*, Cm<sup>r</sup> | This study |
| QT6357 | QT598 $\Delta plfX$::Tn7T-Cm:: *Pplf*$\Delta plfB$ promoter<br>*luxCDABE*, Cm<sup>r</sup> | This study |
| QT6358 | QT598 $\Delta plfX$:: Tn7T-Cm:: *Pplf*$\Delta plfX$ $\Delta plfB$<br>promoter *luxCDABE*, Cm<sup>r</sup> | This study |
| **Plasmids** |  |  |
| pIJ514 | pGP-Tn7-Cm::*luxCDABE*; *rbs* Ap<sup>r</sup>, Cm<sup>r</sup> | (42, 50) |
| pIJ610 | pGp-Tn7-Cm-*lux*:: *Pplf* promoter | This study |
| pIJ621 | pGP-Tn7-Gm::<br>*plf* operon |  |
| pIJ643 | pGp-Tn7-Cm-*lux* :: *Pplf* promoter $\Delta plfB$ Cm<sup>r</sup> | This study |
| pIJ647 | pGp-Tn7-Cm-*lux*:: *Pplf* promoter $\Delta plfX$ Cm<sup>r</sup> | This study |
| pIJ652 | pBlueScript:: *Pplf*$\Delta plfX$<sub>QT598</sub>; Amp<sup>r</sup> | This study |
| pIJ655 | pGp-Tn7-Cm-*lux* :: *Pplf* promoter $\Delta plfX$ $\Delta plfB$ Cm<sup>r</sup> | This study |
| pIJ656 | pMEG375:: *orfE*-Gm<sup>R</sup>-*Pplf*$\Delta plfX$; Amp<sup>r</sup>, Cm<sup>r</sup>, Gm<sup>r</sup> | This study |
| pIJ659 | pET28a-TEV in S17:: *plfB* | This study |
| pIJ661 | pMEG375:: *orfE*-Gm<sup>r</sup>-*PplfΔplfB*; Amp<sup>r</sup>, Cm<sup>r</sup>, Gm<sup>r</sup> | This study |
| pIJ666 | PGP-tn7-CM:: *plfX* | This study |
| pIJ667 | PGP-tn7-CM:: *plfB* | This study |
| pIJ668 | pET28a-TEV in S17:: *plfX* | This study |
| pCP20 | Temperature-sensitive FLP recombinase plasmid;<br>Cm<sup>r</sup>, Amp<sup>r</sup> | (42) |
| pKD4 | Plasmid used for the amplification of the<br>*kan* resistance cassette | (42) |
| pSTNSK | Tn7 Transposase plasmid; pST6K::*tnsABCD*;<br><u>oriSC101(Ts)</u>; Tp<sup>r</sup> | This study |
| pGp-Tn7-Gm | Cloning vector for *att* Tn7 complementation | (42) |
| pGp-Tn7-Cm | Cloning vector for *att* Tn7 complementation | (42) |
| pUCM-T | Cloning-vector; Amp<sup>r</sup> | Bio Basic Inc., Markham,<br>ON., Canada |
| pMEG375 | *sacRB mobRP4 oriR6K*; Cm<sup>r</sup>, Amp<sup>r</sup> | Megan Health, St. Louis,<br>MO |
| pFGM1 | Template plasmid for the FRT-flanked gentamycin<br>resistance cassette; Gm<sup>r</sup>, Amp<sup>r</sup> | (51) |

### Construction of the *plf-lux* reporter fusion

Transcriptional fusions were generated using Tn7 transposition, in which the *plf* promoter region and segments of the predicted *plf* regulatory region were fused to the *lux* operon. The whole promoter region and putative internal regulators were amplified using primers presented in Table 2. The amplified fragments were ligated into the luciferase encoding vector pIJ514 digested with *XhoI* and *SpeI* to construct plasmids pIJ610, pIJ643, and pIJ647, respectively. Plasmid pIJ610 contains the entire *plf* promoter region spanning from predicted promoter 6 (P6) to the end of predicted promoter 4 (P4), whereas pIJ643 and pIJ647 contain promoter fragments lacking the *plfB* or *plfX* encoding regions, respectively. The ligation mixtures were purified and subsequently transformed into *E. coli* DH5α λpir, followed by selection on LB plates containing chloramphenicol. Plasmids pIJ610, pIJ643, and pIJ647 were transformed into *E. coli* SM10 λpir-derived strain MGN-617 which was used as the donor for conjugative transfer. The recipient strains were strains QT598 and QT598Δ*plf* containing the plasmid pSTNSK expressing the Tn7 *tnsABCD* transposase genes. Conjugation was carried out overnight on LB agar plates supplemented with diaminopimelic acid (DAP) 0.3 mM to permit growth of the donor strain and at 30°C to maintain the temperature sensitive Tn7 tranposase encoding plasmid. Following conjugation overnight, bacteria were then suspended in 1 ml of phosphate-buffered saline (PBS), washed twice in PBS pH 7.4, serially diluted, and cultured on LB agar supplemented with chloramphenicol, and incubated at 37°C. The integration at the *att*Tn7 site and loss of the transposase-encoding plasmid pSTNSK in chloramphenicol-resistant colonies was verified by confirming sensitivity to kanamycin and ampicillin. Finally, the insertions at the *att*Tn7 site were confirmed by PCR using primers CMD26 and CMD1416, listed in Table 2.

**TABLE 2.**
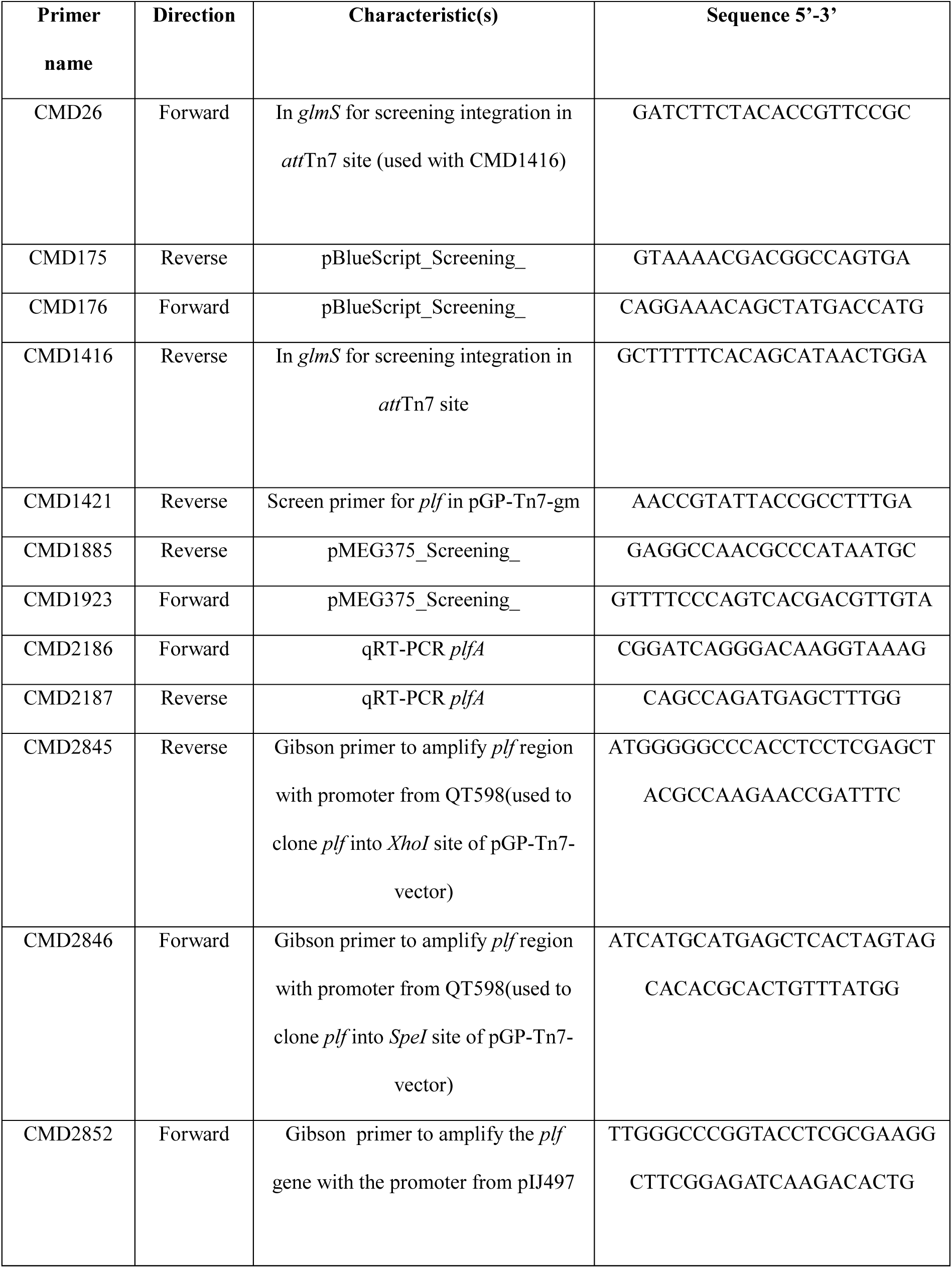
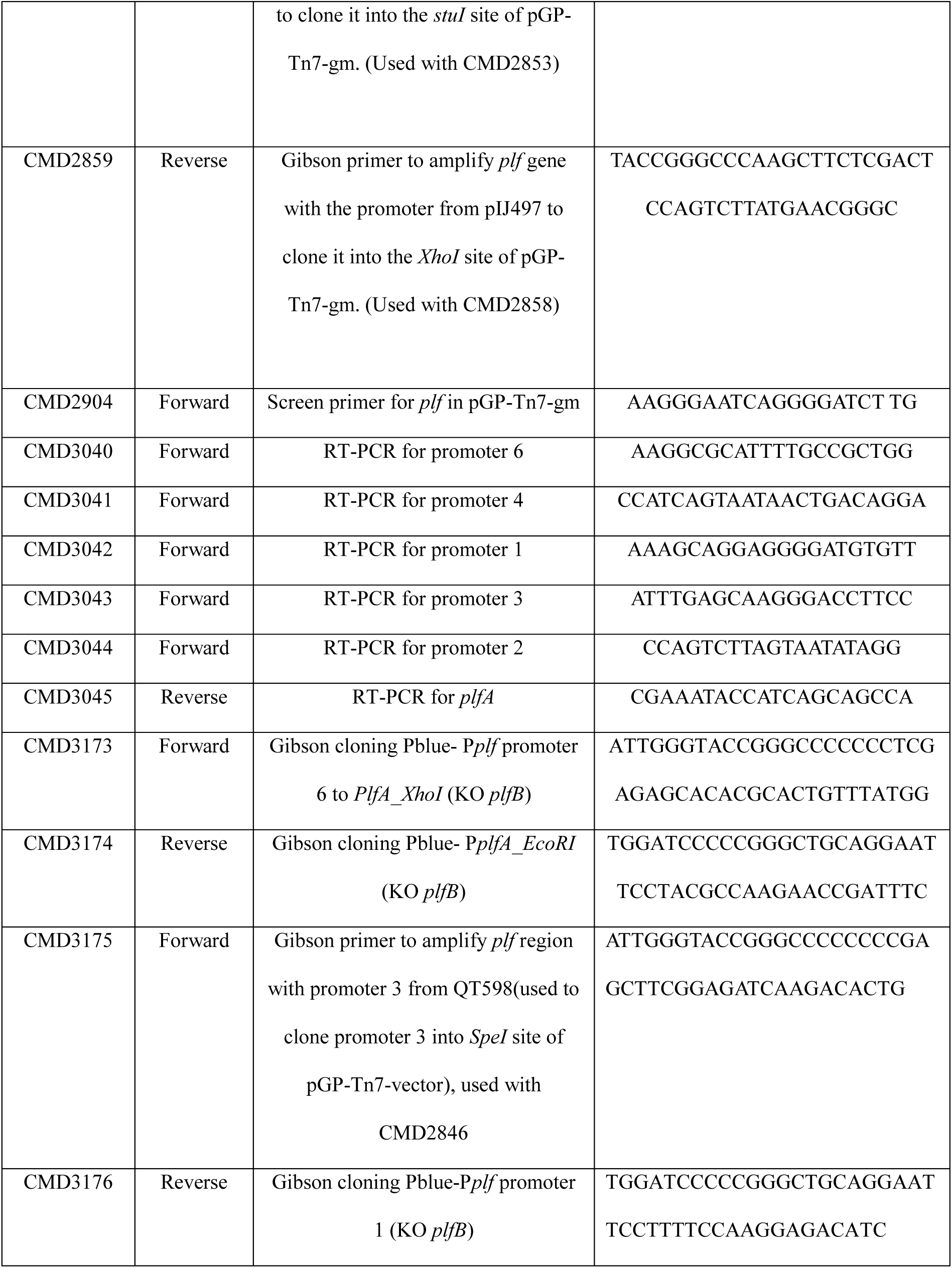
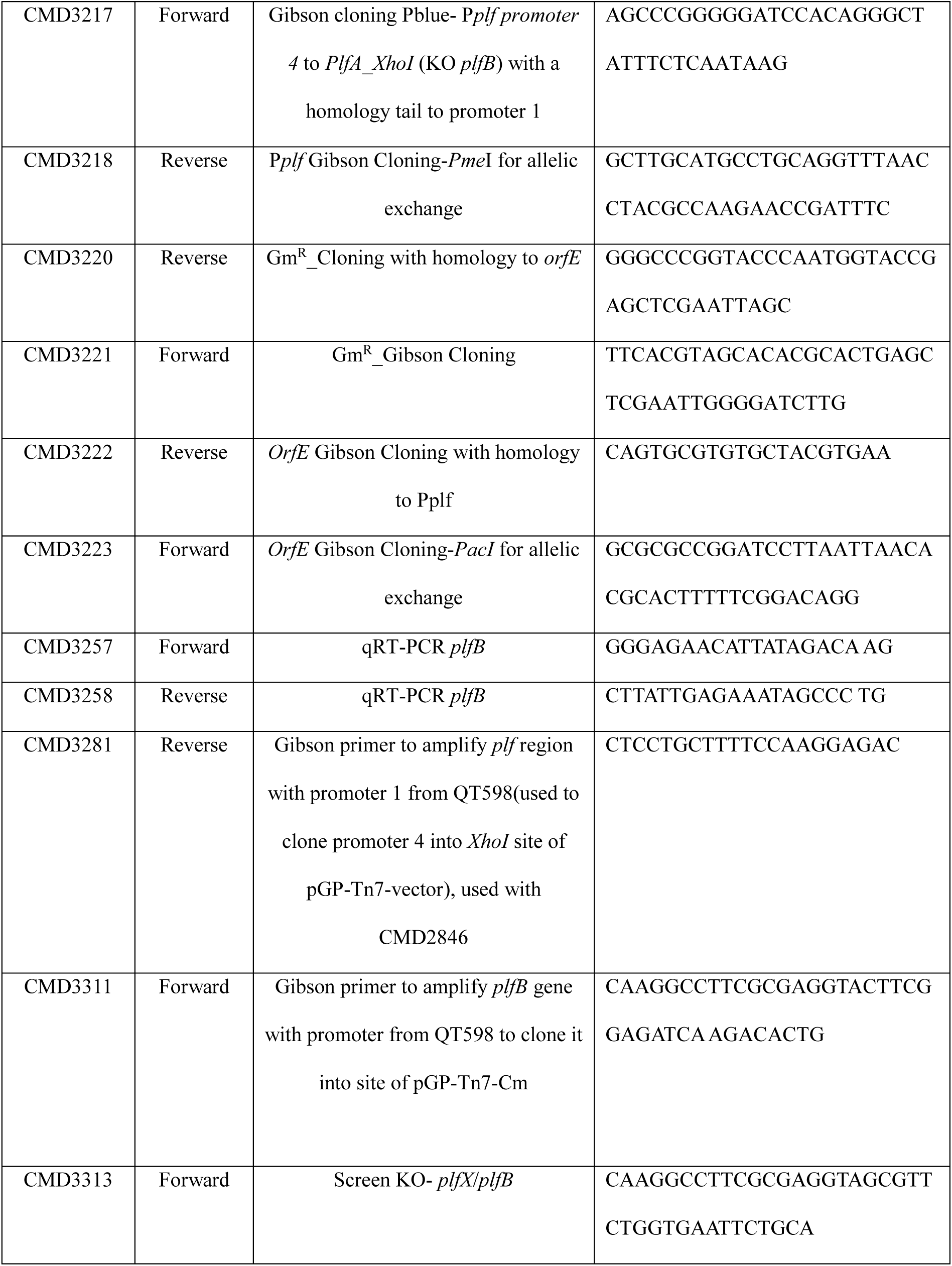
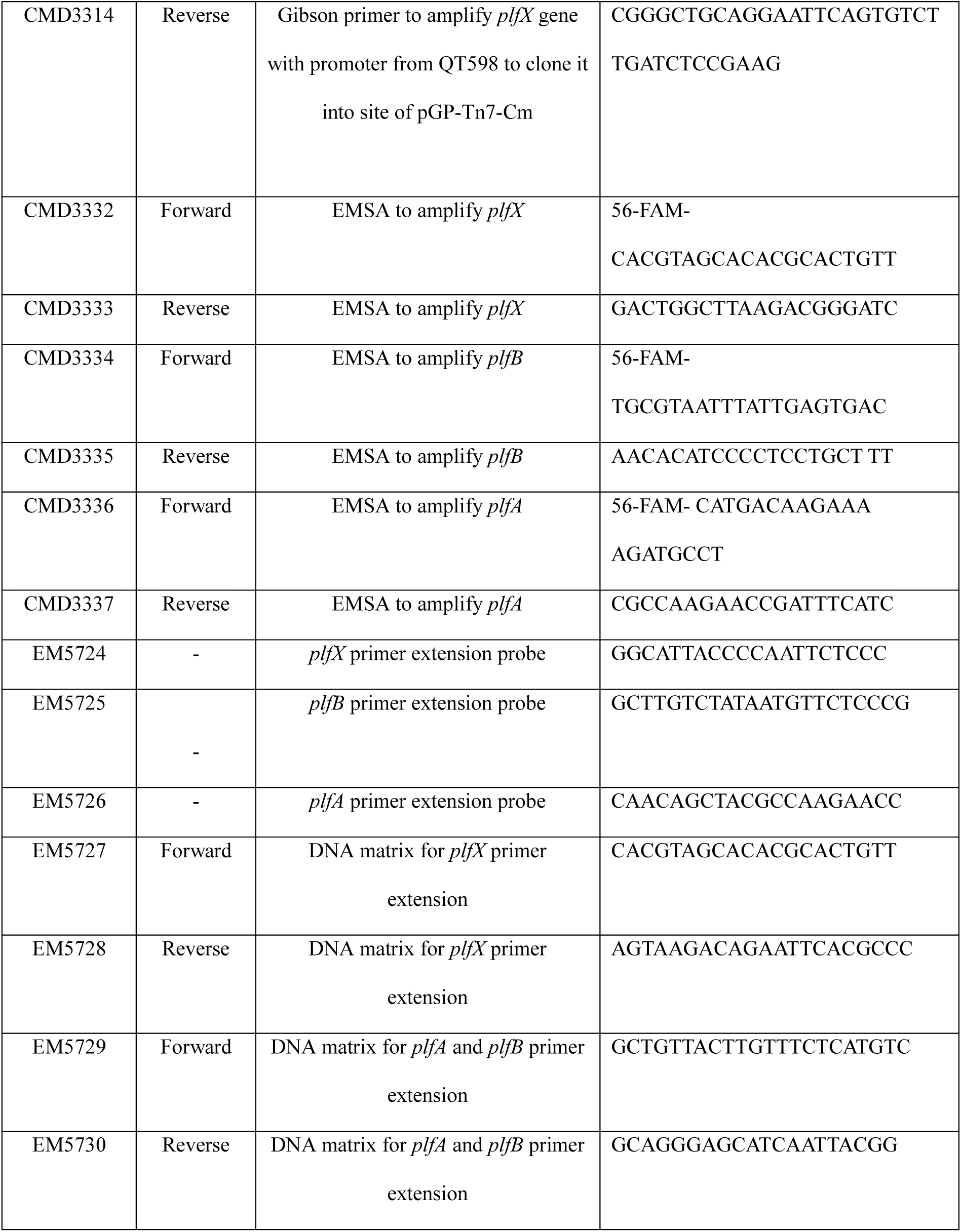
List of primers.

### Deletion of genes from strain QT598 and complementation

The *plf* gene deletion mutants, and deletion mutants for some unknown regulatory genes such as Δ*hns* and Δ*argR* mutants were generated by the Datsenko and Wanner method using plasmid pKD4 as a template to amplify the kanamycin resistance cassette (41). The primers used to generate mutants are listed in Table 2. Plasmid pCP20 expressing the FLP recombinase was transformed into the mutant strains to remove the antibiotic resistance cassette flanked by the FLP recombination target (FRT). The deletion of possible internal regulators was done by the allelic exchange method which is explained in another section. Cloning of *plfB and plfX* genes was performed by PCR amplification of DNA segments including the native promoter regions of each of the genes from genomic DNA of strain QT598, using specific primers (Table 2) and Phusion flash high-fidelity PCR master mix (Thermo Fisher Scientific, Waltham, MA, USA). The inserts obtained from PCR amplification contained Gibson homologies to the multi-cloning site (MCS) of the mini-Tn7 containing plasmids specific to the *KpnI* and *EcoRI* restriction sites of pGP_miniTn7_CM. DNA segments were assembled using the pEASY-UNI assembly kit (TransGen Biotech, Beijing, China) to pGP_miniTn7_Cm linearized by *KpnI* and *EcoRI* (New England Biolabs, Ipswich, MA, USA) to obtain plasmids pIJ666 and pIJ667.

### Testing the *Pplf-lux* fusions under different conditions

To investigate the effects of different growth conditions on expression from the *plf* promoter, different media and growth conditions were used. Media included Nutrient agar (Alpha Bioscience, Baltimore, MD, USA), MacConkey-lactose agar (Alpha Bioscience, Baltimore, MD, USA), Brain Heart Infusion (BHI) (Alpha Bioscience, Baltimore, MD, USA), LB with and without shaking and LB agar (LBA), and LBA buffered with 0.1 M Na_2_HPO_4_-NaH_2_PO_4_ (42), LBA with multiple concentrations of NaCl, M9 minimal media with different carbon sources including glucose (C_6_H_12_O_6_) (0.20% w/v) or glycerol (C_3_H_8_O_3_) (0.20%) (43). Overnight cultures of strains containing p*plf-lux* fusions were cultured with agitation at 250 rpm in 5 ml LB medium or on agar plates overnight. The next day, 3 μl of each broth overnight culture was transferred to 180 μL of broth media and incubated with agitation until mid-logarithmic phase and stationary phase had been reached in 96-well plates (Corning white with clear flat bottom). For cultures maintained on agar, colonies were passaged three times, scraped, and resuspended. Then serial dilutions were prepared to obtain an OD₆₀₀ (∼1) equivalent to ∼8 × 10⁸ CFU·mL⁻¹. Volumes of 200 μL of these samples were then added to 96-well plates (Corning white with clear flat bottom). The luminescence was measured using a Cytation™ 3 Cell Imaging Multi-Mode Reader (BioTek Instruments Inc.). The luminescence results were reported as relative luminescence units (RLU). The luminescence readings were normalized to the O.D. _600nm_ values. The luminescence of *lux* fusion-containing strains streaked on LB agar plates was also measured with a ChemiDoc XRS system equipped with Quantity One 1-D analysis software (Syngene Chemi Genius).

### RT-PCR

Total RNA was extracted from bacterial cultures grown in biological triplicates from 3 successive passages on LBA and immediately stabilized after extraction. The RNA was treated with Turbo DNase (Ambion, Thermo Fisher Scientific, St. Laurent, QC, Canada) and then checked for quality and integrity using a Nanodrop (ND-1000) prior to cDNA synthesis. First-strand cDNA was synthesized from purified RNA using standard reverse transcription reagents and the Advanced 1-Step RT-PCR Kit (Wisent, St. Bruno, QC, Canada). Detectable transcripts from predicted promoters were determined by PCR from the cDNA samples. The primers are listed in Table 2. PCR products were analyzed by agarose gel electrophoresis and visualized under UV light. Transcriptional activity from predicted promoter regions was assessed by end-point RT-PCR, with the aim of detecting the presence or absence of specific transcripts rather than performing quantitative expression analysis. Therefore, no housekeeping or constitutively expressed genes were used for normalization. PCR products were analyzed by agarose gel electrophoresis and visualized under UV illumination. Primers used in this study are listed in Table 2.

### Primer extension

Total RNA (20µg) from bacterial cultures was extracted, following growth on LBA for 3 successive passages, and then treated with DNase to remove contamination. Primer extension was used to determine each gene’s transcriptional start point (Table 2). A radiolabelled oligonucleotide complementary to the mRNA downstream of the promoter was annealed to the RNA and extended with reverse transcriptase to the 5′ end of the transcript. Alongside a sequencing ladder made from the matching DNA template, the resultant cDNA fragments were sorted on a denaturing 8% polyacrylamide gel and then revealed with a GE Healthcare Typhoon Trio (15). Reverse transcriptase-deficient negative controls confirmed that there was no DNA contamination.

### Allelic Exchange to knock out the *plfX* and *plfB* genes

To construct the QT598***Δ****plfX* mutant, the desired fragment was amplified using primers CMD3175 and CMD3174 from QT598 by PCR using Phusion Flash High-Fidelity Master Mix (Thermo Fisher Scientific, Waltham, MA, USA) and cloned into pBlueScript II SK+ (Agilent Technologies), digested with *XhoI and EcoRI* (NEB) to generate pIJ652. For QT598***Δ****plfB,* the desired insert was assembled by ligating the fragment amplified with CMD3173 and CMD3176 to the fragment amplified with CMD3217 and CMD3174, after which the combined product was cloned into *XhoI/EcoRI*-digested pBlueScript via Gibson assembly using NEBuilder HiFi DNA Assembly Master Mix (NEB, Ipswich, MA, USA) to generate pIJ672. These constructs were verified by PCR using primers CMD175 and CMD176, which flank the pBlueScript multiple cloning site. The *PplfΔplfX* region was then amplified from pIJ652 using CMD3175 and CMD3218, and the *PplfΔplfB* region was amplified from pIJ672 using CMD3173 and CMD3218. These fragments were subcloned into suicide vector pMEG375, digested with *PacI* and *PmeI* (NEB, Ipswich, MA, USA), together with the *orfE* fragment (the region adjacent to *plfX* on pEC598, used for homologous recombination) amplified from QT598 using CMD3223 and CMD3222, then the Gm cassette amplified from pFGM1 using CMD3221 and CMD3220. This assembly generated the allelic-exchange vectors pMEG375::*orfE-Gm^R^-PplfΔplfX* (pIJ656) and pMEG375::*orfE-Gm^R^-PplfΔplfB* (pIJ661). Final constructs were verified by PCR using screening primers CMD1923 and CMD1885, which flank the pMEG375 multiple cloning site. The knockout mutations of each of these genes were generated by allelic exchange as described previously (44). Allelic exchange vectors carrying fragments of *plfX* and *plfB* (pIJ656 and pIJ661) were first transferred from the *E. coli* χ7213 donor strain to QT598 by conjugation. Transconjugants carrying single-crossover plasmid insertions in the *Pplf* promoter region, which lies between *plfA* and the upstream *orfE* locus, were selected on LB agar containing gentamicin but lacking DAP. Double-crossover recombinants, which result in the loss of the allelic exchange vector, were subsequently isolated by *sacB*-mediated counterselection on LB agar containing gentamicin and 5% sucrose, without NaCl (45, 46). The FRT-flanked gentamicin resistance cassettes were then excised by transforming the recombinant strains with the FLP recombinase-expressing, temperature-sensitive plasmid pCP20. The resulting double-crossover knockout mutants QT598Δ*plfX* and QT598Δ*plfB* (QT6349 and QT6308) were confirmed by screening for antibiotic sensitivity and by PCR using primers CMD3313 and CMD3045.

### Protein Preparation and Western Blot Analysis

Preparation of fimbrial extracts and western blotting were performed as described previously with some modifications (6). The overnight cultures on LBA plates were scraped, resuspended in 1 ml PBS and then incubated at 55 °C for 1 h. Supernatants were then centrifuged at 6000 x *g* for 15 minutes, and the aliquots were precipitated in 15% trichloroacetic acid (TCA) (v/v). Subsequently, the samples were incubated at 4 °C for 1 h with shaking. Followed by centrifugation at 14,000 × *g* for 20 min at 4°C, the pellets were washed twice with 0.5 M Tris-HCl-0.5 M EDTA (pH 12.0) and resuspended in 0.5 M Tris-EDTA. The samples were migrated and resolved on 15% SDS-PAGE gels at 200 V for 40 min, after the loading dye, 4x Laemmli sample buffer (Tris-HCL 200 mM, 8% SDS (v/v), 40% glycerol (v/v), 4% β-mercaptoethanol (v/v), 50 mM EDTA, and 0.4% bromophenol blue (v/v) was added to them. Protein samples were then transferred onto a nitrocellulose membrane for 60 min at 100 V (Pall Corporation, Port Washington, USA). Then the membrane was blocked by using tris-buffered saline (TBS) with 0.1% Tween-20 (TBST), and 5% (w/v) skim milk for 1 h at room temperature with shaking. Incubations with rabbit polyclonal antibodies provided by New England Peptide (1:1,000) against a peptide corresponding to part of the PlfA major fimbrial subunit (6) and secondary goat anti-rabbit (1:20,000) antibodies were carried out for 1 h at room temperature. Signals were then revealed using SuperSignal™ West Pico PLUS Chemiluminescent Substrate (Thermo Fisher Scientific, Waltham, MA, USA) and visualized with the ChemiGenius 2 documentation system according to the manufacturer’s instructions.

### Hemagglutination assays

Macro-Hemagglutination tests were performed in the presence of 2.5% D-mannose to block agglutination due to type 1 fimbriae. Turkey and human O⁺ erythrocytes were washed twice with PBS, centrifuged, and resuspended in PBS to a final concentration of 3%. Strains such as QT5726 (clone producing PL fimbriae with PlfG class II adhesin from QT598), QT598, QT6308 (QT598Δ*plfB)*, and QT6335 (QT6308 complemented with *plfB)*, and QT6349 (QT598Δ*plfX)*, were grown on LB agar until the 3^rd^ passage at 37 °C, then scraped and washed with PBS, and centrifuged at 3,000 × *g* for 15 min, suspended in PBS (pH 7.4) and adjusted to an optical density at 600 nm (OD_600_) of 0.6 (47, 48).

### Turkey infection model and tissue sample collection

The animal study was reviewed and approved by the animal ethics evaluation committee-Comité Institutionel de Protection des Animaux (CIPA project No. 2311–01) of INRS. The turkey infections were carried out as described previously. The 6-day-old poults were injected in the left caudal air sac with either the wild-type *E. coli* strain QT598, the QT598 Δ*plfB* mutant, or the complemented *plfB* strain (QT6335) with 2.8 × 10^8^ CFU in 100 µL of inoculum. After 48 h, poults were euthanized; lungs, spleens, and kidneys were sampled, homogenized, diluted, and plated on MacConkey agar for enumeration of bacterial colonies.

### RNA extraction for quantification of gene expression *in vivo* and *in vitro*

Bacterial cultures were grown in triplicate in LBA. RNAprotect (Qiagen, Toronto, ON, Canada) was added to cultures and RNA extracted using an EZ-10 Spin Column Total RNA Miniprep Kit (BioBasic) according to the manufacturer’s instructions. Ambion Turbo DNase (Thermo Fisher Scientific, St. Laurent, QC, Canada) was used to remove contaminating DNA. RNA integrity was assessed using a Nanodrop (ND-1000) (Thermo Fisher Scientific, St. Laurent, QC, Canada). RNA was also extracted from infected lung tissues at 48 h p.i. with TRIzol reagent (Thermo Fisher Scientific, St. Laurent, QC, Canada), followed by DNase 1 treatment. Total RNAs were then reverse-transcribed to cDNAs using *TransScript*® all-in-one First-Strand cDNA Synthesis SuperMix Kit (TransGen, Haidian District, Beijing, China). The *rpoD* gene was used as a housekeeping control. Each qRT-PCR run was done in triplicate, and for each reaction, the calculated threshold cycle (*C_T_*) was normalized to the *C_T_* of the *rpoD* gene amplified from the corresponding sample. The foldchange was calculated using the 2^−ΔΔ^*CT* method. Primers used for qRT-PCR analysis are listed in Table 2.

### Immunofluorescence to detect PL fimbriae on bacterial cells

Bacterial strains such as ORN172 PlfG II (QT5726), QT598, and QT598Δ*plf* (QT5973) were grown in LB broth for overnight or on LBA or LBA 0.3 M NaCl plates for 3 successive passages at 37°C. For each of the samples, bacterial pellets were obtained by centrifugation at 4000 x *g* for 6 min, followed by washing 2 times with PBS and dilution to a final bacterial cell count of ∼1 × 10^⁸^ CFU/mL. Then, bacterial cells were fixed with 2% paraformaldehyde (PFA, E. Merck AG, Darmstadt, Federal Republic of Germany) for 45 min at 4°C and subsequently washed with PBS 3 times and then incubated with polyclonal antibody (1:1,000) against the whole PL fimbriae for 45 min at 4°C on a shaker. After that, samples were washed 2 times with PBS, and they were incubated with secondary antibody Alexa Fluor*™* 488 (Thermofisher, USA, 5 µg/ml in PBS, excitation 490, emission 525 nm) (1:1,0000), in PBS for 45 min at 4°C on a shaker and washed three times using PBS.

Finally, 3 µL of each sample were transferred to microscope slides, separately. The nuclei of bacteria were then stained with 4′,6-diamidino-2-phenylindole (DAPI) 1µg/ml (Thermofisher, USA, excitation 358 nm, emission 461 nm) for 5 min at room temperature in the dark, followed by 3 washing steps with PBS. A coverslip was placed on the slides for visualization by confocal microscopy. Confocal images were acquired using a Zeiss LSM 780 microscope (Carl Zeiss, Germany) with sequential laser excitation for DAPI, Alexa Fluor 555, and GFP. Z-stack images were collected through the epithelial surface of OCT-embedded lung tissue. ZEN Black software (version 2012 Sp4; Zeiss) was used for image acquisition, channel separation, and intensity calibration.

### Protein expression and purification

*E. coli* BL21(DE3) was used as the bacterial host for the expression vectors. The expression vectors encoding PlfB or PlfX were prepared by the manufacturer (GenScript, Piscataway, NJ, USA) with the His-tag placed on the N-terminal. Strains were grown on Terrific Broth (TB) at 37°C overnight, and then an initial 12 mL bacterial culture was transferred into 2 L of fresh growth medium at 37°C until OD₆₀₀ ≈ 0.5–0.6. This was followed by incubating overnight at 25°C after adding Isopropyl β-D-1-thiogalactopyranoside (IPTG) (Sigma Aldrich, St. Louis, MO, USA) to final concentrations of 0.1, 0.5, or 1 mM. The soluble and insoluble fractions were analyzed by SDS-PAGE to determine the localization of the recombinant protein after harvesting and lysing of the cells in a small-scale expression (49). The large-scale expression was performed under the desired conditions, and the cell pellets were resuspended in lysis buffer (50 mM Tris-HCl, 300 mM NaCl, 10 mM imidazole, pH 8.0). Next, centrifugation at 15,000 *× g* for 20 min at 4 °C was used to clarify the lysates. Bio-Rad gravity-flow columns filled with HisPur™ Ni-NTA resin (Thermo Fisher Scientific) were used for the purification, and after that, the resins were washed, and the bound protein was eluted by various concentrations of imidazole ranging from 50 to 500 mM. All the fractions were analyzed by SDS-PAGE and prepared using the 4x Laemmli sample buffer (Tris-HCL 200 mM, 8% SDS (v/v), 40% glycerol (v/v), 4% β-mercaptoethanol (v/v), 50 mM EDTA, and 0.4% bromophenol blue (v/v)) and resolved in 10% SDS-PAGE gel at 200 V for 40 min and stained with Coomassie blue to visualize protein bands. The Bradford assay (Bio-Rad) with bovine serum albumin (BSA) standards was used to determine the protein concentrations.

### Electrophoretic mobility shift assay (EMSA)

For EMSA experiments, PlfB and PlfX were expressed as recombinant proteins using constructs containing the complete coding sequences of plfB and plfX, respectively. The corresponding expression constructs and recombinant protein sequences used for purification are shown in Fig. S7. Recombinant proteins were expressed in pET28a-TEV using BL21 producing [His TAG]-tagged PlfB and PlfX proteins. Proteins were purified, and then purified proteins in increasing concentrations were incubated with a fixed amount of labelled DNA fragments (containing each *plf* promoter region ranging from 100–300 bp) in binding buffer (10 mM Tris-HCl, pH 7.5, 50 mM KCl, 1 mM DTT, 10% glycerol) at room temperature for 20 minutes. Then the EMSA loading dye (50mM Tris-HCl, pH 7.4, 50mM EDTA, 0.1% bromophenol blue, 12.5% glycerol) was added to each sample, and samples were resolved on a 6-8% native polyacrylamide gel in 1×TAE buffer. DNA–protein complexes were visualized by Typhoon™ FLA 9000 biomolecular imager (GE Healthcare/Cytiva, Marlborough, MA, USA).

### Statistical Analyses

Data are shown as mean ± SEM. Two-group comparisons used a two-tailed Student’s t-test, while multiple groups were analyzed by one-way ANOVA. For turkey infection studies, the Mann-Whitney U test or Kruskal-Wallis test was applied. P < 0.05 was considered significant. Analyses were performed with GraphPad Prism v7 (GraphPad Software, San Diego, CA, USA).

## Ethics statement

All animal and blood procedures were approved by the Institutional Animal Care and Use Committee (CIPA) and the Institutional Committee on Research Ethics (CER) of INRS-Armand-Frappier Santé Biotechnologie Research Centre. Turkey air sac infection and blood collection protocols were conducted under CIPA protocol 2311-01 and CER protocol 19-507, respectively, following institutional and national ethical guidelines.

## Data Availability

All relevant data are within the manuscript and its Supporting Information files.

## Acknowledgments

We thank Jessy Tremblay for assistance with confocal microscopy and Prof. Fredric Veyrier and Francisco Pulido for assistance with the genome sequencing, and Marlène Fortier for protein extraction and purification.

## Author Contributions

Conceptualization: Fariba Akrami, Charles M. Dozois.

Data curation: Fariba Akrami.

Formal analysis: Fariba Akrami, Tylor Pereira, Hossein Jamali, Sebastien Houle, Eric Masse,

Evelyne Ng Kwan Lim, France Daigle, Charles M. Dozois.

Funding acquisition: Charles M. Dozois.

Investigation: Fariba Akrami.

Methodology: Fariba Akrami, Tylor Pereira, Hossein Jamali, Sebastien Houle, Eric Massé,

Evelyne Ng Kwan Lim, France Daigle, Charles M. Dozois

Project administration: Fariba Akrami, Eric Masse, France Daigle, Charles M. Dozois.

Resource: Charles M. Dozois.

Supervision: Charles M. Dozois and France Daigel.

Validation: Fariba Akrami, Eric Masse, France Daigle, Charles M. Dozois.

Writing-original draft: Fariba Akrami and Charles M. Dozois.

Writing– review & editing: Fariba Akrami, Eric Massé, France Daigle, Charles M. Dozois.

## Supplementary data

## Supplementary data

**FIG. S1.**
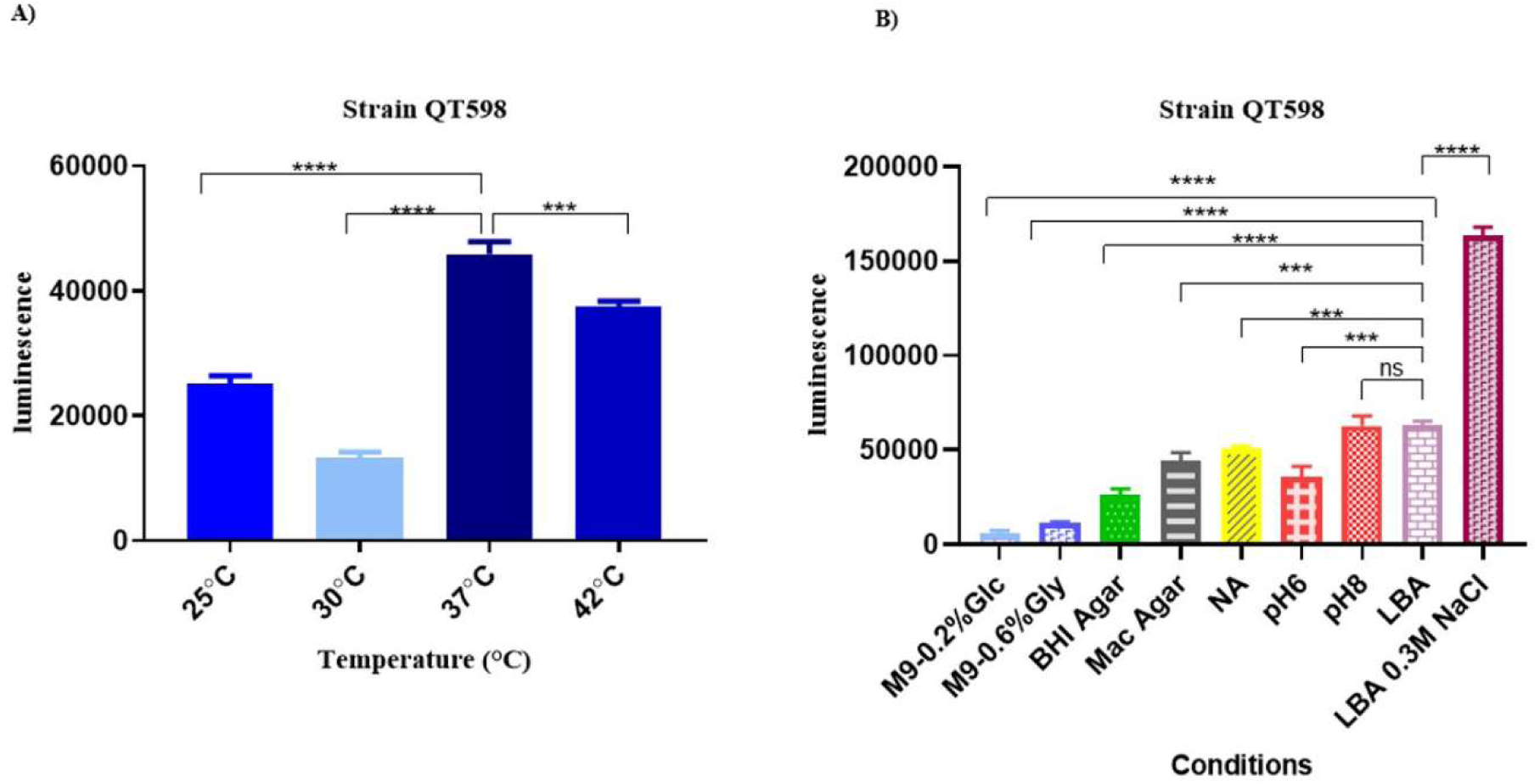
Luminescence levels of strain QT598 p*plf-lux* under different growth conditions. **A)** QT598 P*plf-lux* was grown for 3 successive passages in LBA at different temperature. **B)** QT598 P*plf-lux* was grown for 3 successive passages in various media, including minimal M9 with glucose and glycerol, BHI agar, MacConkey agar, nutrient agar, pH-adjusted media, Luria-Bertani (LBA), and LBA with 0.3 M NaCl. Luminescence, reflecting bacterial activity, was measured for each condition. Data are the means from three independent experiments, and error bars represent standard errors of the means: ns= not significant *** *P* < 0.001 and **** *P* < 0.0001 using one-way ANOVA.

**FIG. S2.**
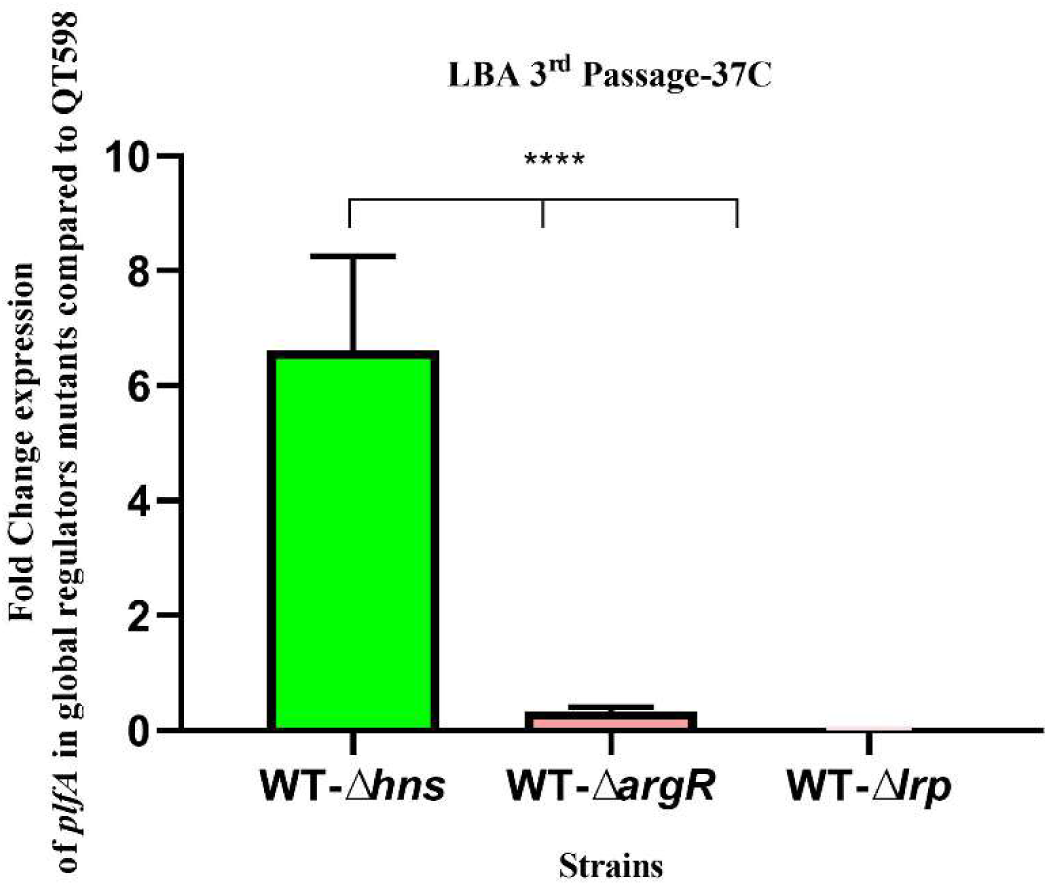
Differential expression of *plfA* in mutants compared to wild type in LBA. Quantitative RT-PCR analysis showing the fold change in *plfA* expression in global regulator mutants compared to QT598 grown on LB plates. Expression of *plfA* was significantly higher in the QT598 Δ*hns* compared to the QT598. Data represent the mean ± standard errors of the mean (SEM) from biological replicates. Data are the means from three independent experiments, and error bars represent standard errors of the means. *, *P* < 0.05; **, *P* < 0.01; and ***, *P* < 0.001, ****, *P* < 0.0001 using one-way ANOVA.

**FIG. S3.**
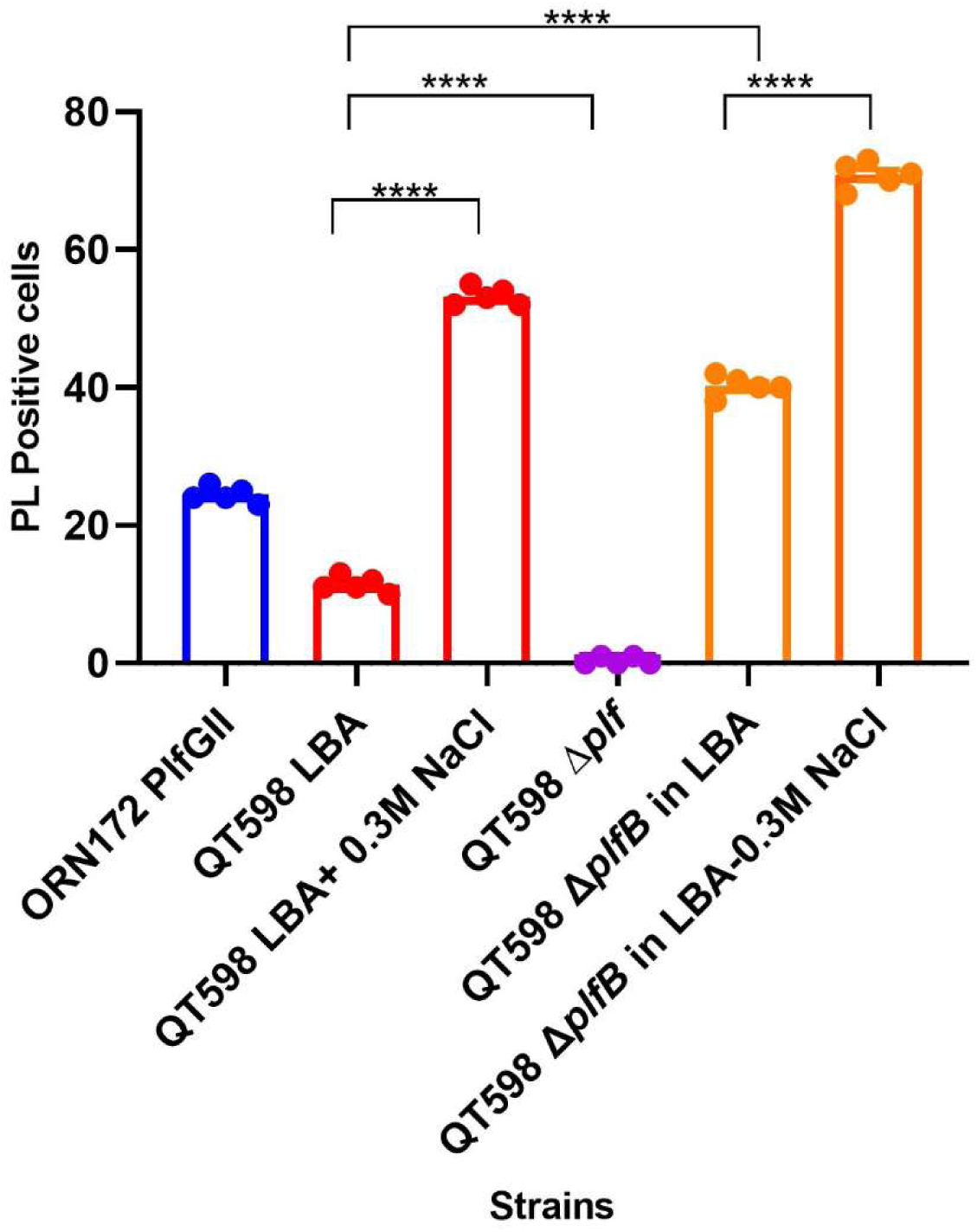
Quantification of PL fimbriae expression by confocal microscopy. PL fimbriae expression was quantified by manually counting PL-positive bacterial cells across multiple, independently selected microscopic fields. Each data point corresponds to one replicate, representing approximately 1,000 cells analyzed from 10 randomly chosen fields. Experiments were conducted on three separate days, with three replicates per condition. Bars represent the mean ± SD.

**FIG. S4.**
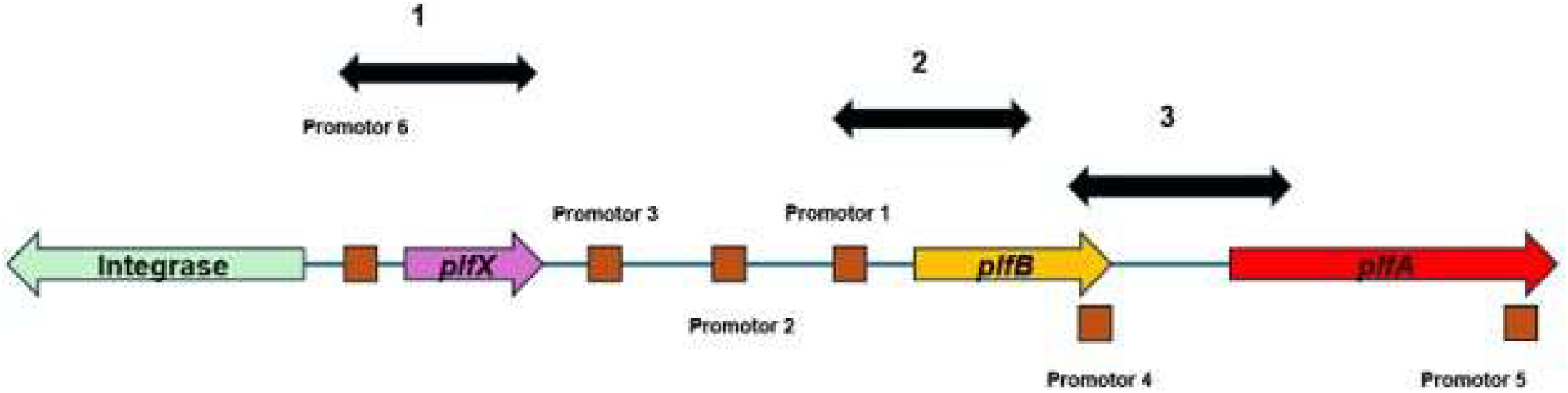
Detection of transcripts from predicted promoters by RT-PCR. Transcripts from predicted promoters were detected by PCR on the cDNA samples. The number 1 is promoter 6 before *plfX*, the number 2 is promoter 1 for *plfB*, and the number 3 is promoter 4 for *plfA*. No products were detected for putative promoters 3, 2, or 5.

**FIG. S5.**
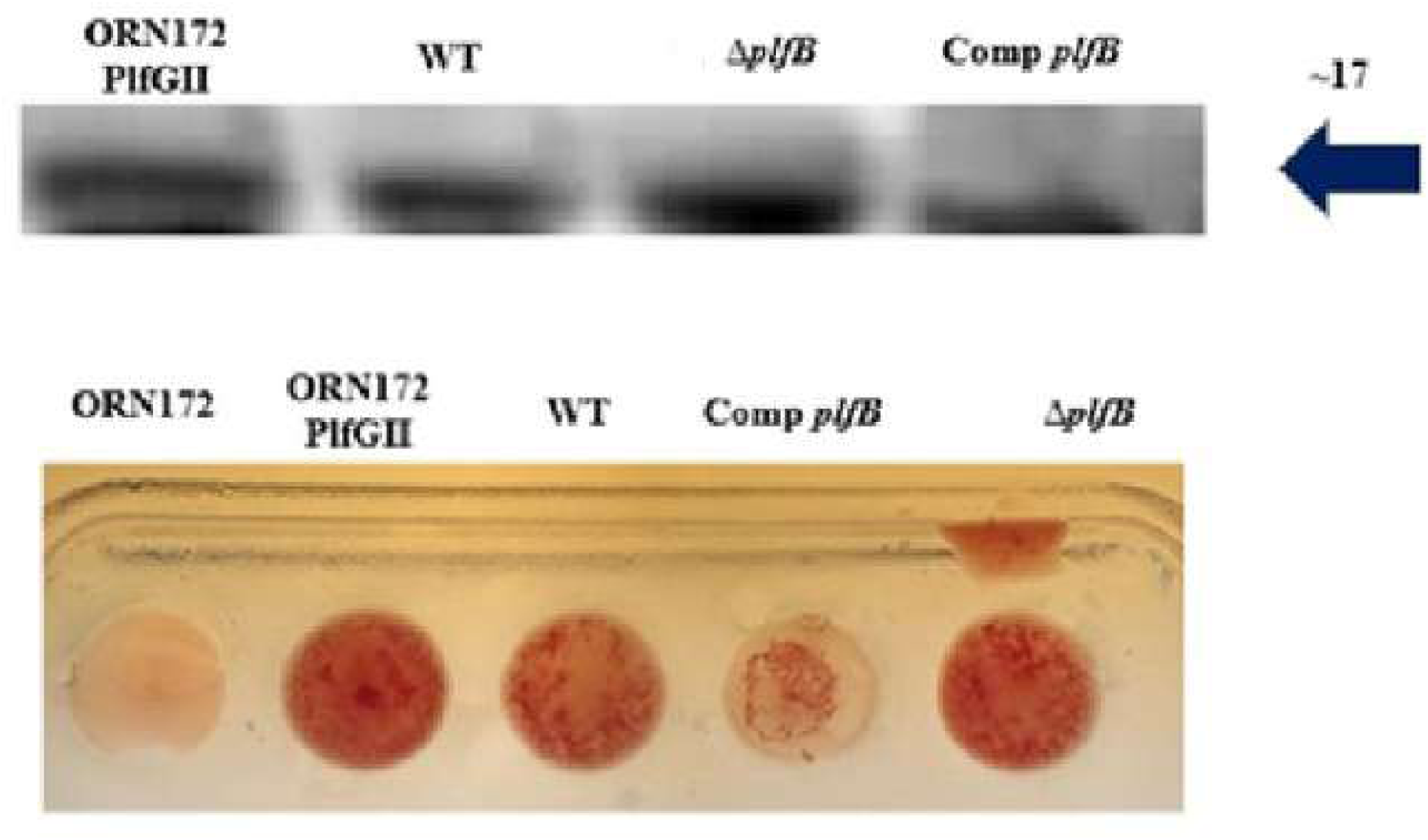
Phenotypic tests for confirmation of *plfB*. Western blot for PlfA demonstrated these trends: stronger in QT598Δ*plfB*, and no difference in Comp *plfB* compared to QT598 (upper panel). Macro-hemagglutination of human O⁺ erythrocytes with 2.5% D-mannose. There was no difference in hemagglutination between QT598 and Comp *plfB,* but QT598Δ*plfB* showed stronger agglutination (lower panel).

**FIG. S6.**
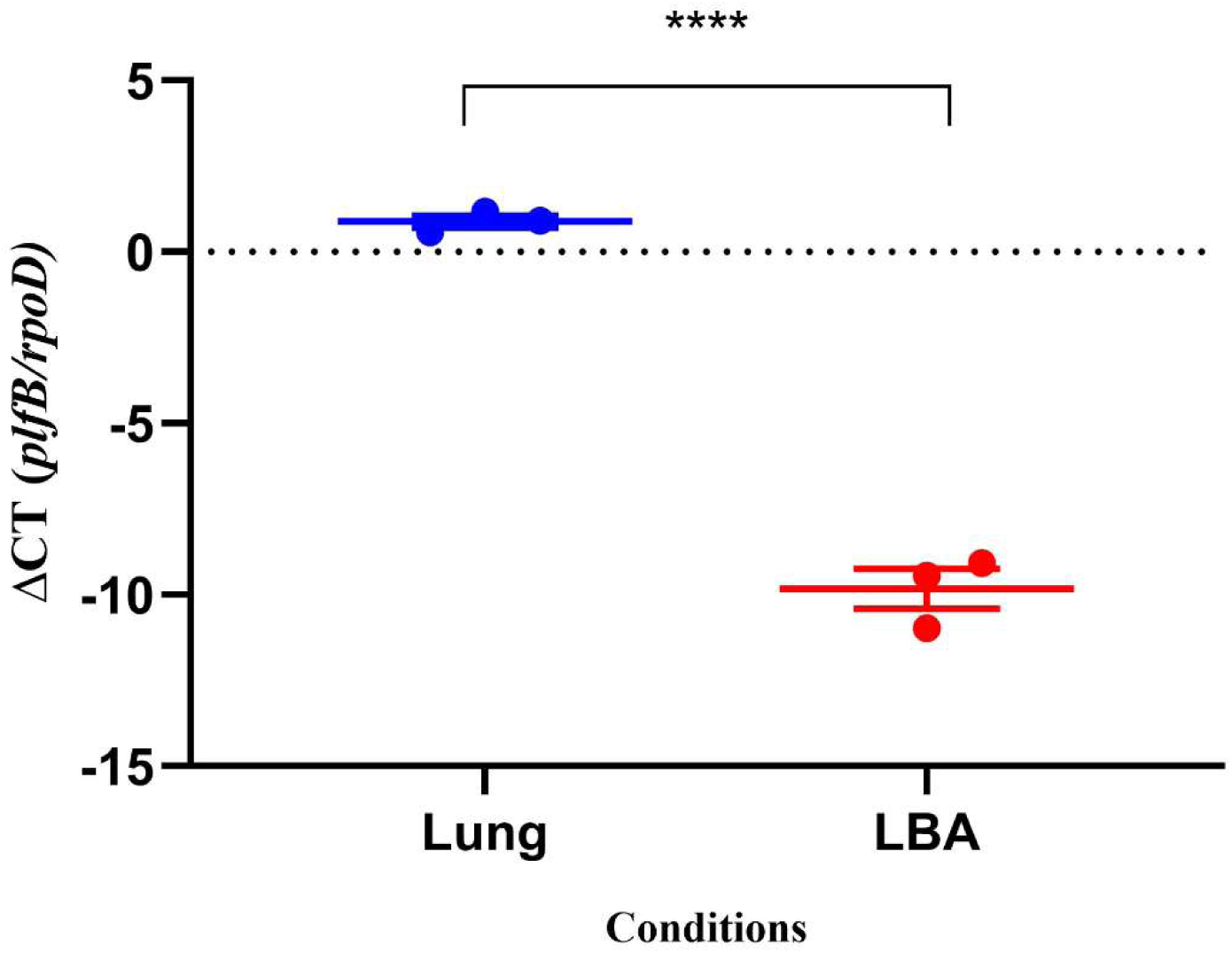
The comparison of *plfB* expression in QT598 *in vivo* to *in vitro*. Quantitative RT-PCR analysis of *plfB* expression was performed using RNA extracted from lung tissues of infected turkeys (Lung) and from bacteria grown on LBA. Transcript levels were normalized to the housekeeping gene *rpoD*, and expression is presented as ΔCt values (Ct*_plfB_* − Ct*_rpoD_*). Each dot represents an individual biological sample, and horizontal bars indicate the mean ± SD. Lower ΔCt values correspond to higher relative expression of *plfB*.

**FIG. S7.**
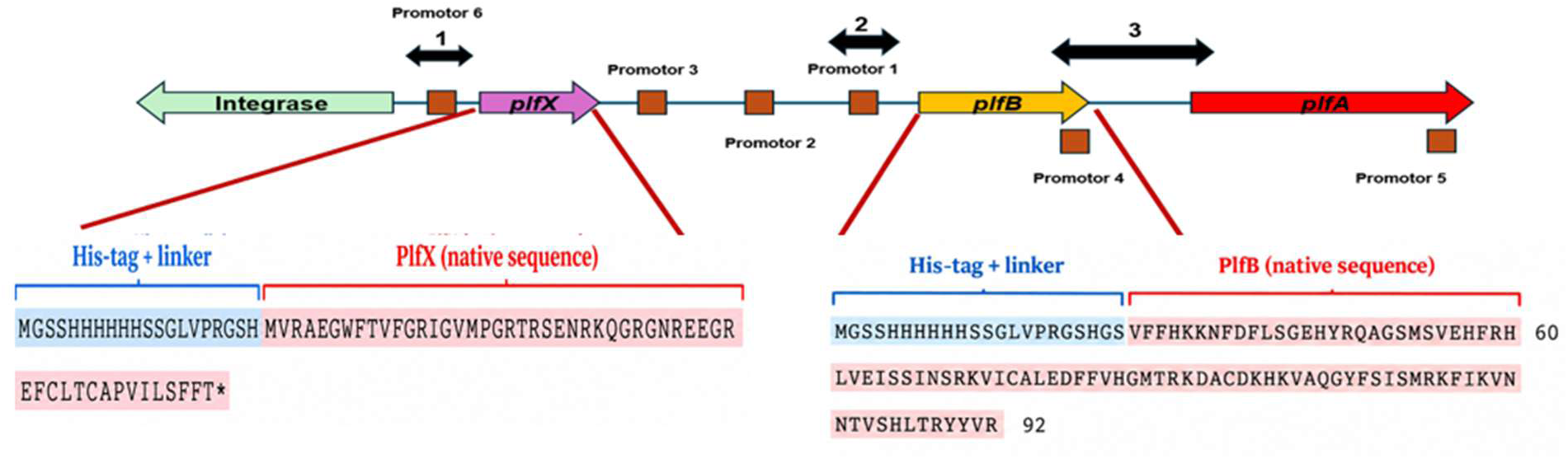
Schematic of DNA fragments used for EMSA. Three separate DNA fragments from the *plf* gene cluster were tested for protein binding. Fragment 1 corresponds to the *plfX* promoter (promoter 6), Fragment 2 to the *plfB* promoter (promoter 1), and Fragment 3 spans the *plfA* promoter region (promoter 4) up to the middle of the gene. The protein sequences of plfB and plfX proteins were indicated below each.

